# Clinical-grade cryopreservation unlocks transplant-ready human pancreatic and stem cell–derived islets for diabetes therapy

**DOI:** 10.64898/2026.04.25.720819

**Authors:** Joseph Sushil Rao, Zongqi Guo, Zenith Khashim, Kyle Knofczynski, Diane Tobolt, Sharath Belame Shivakumar, Parthasarathy Rangarajan, Adam Herman, Muhamad Abdulla, Anna Tran, Benjamin J. Fisher, Tiana Salomon, Sean Lewis-Brinkman, Magie Steinhoff, Appakalai N. Balamurugan, Michael L. Etheridge, Bernhard J. Hering, Sabarinathan Ramachandran, Quinn P. Peterson, John C. Bischof, Erik B. Finger

**Affiliations:** Department of Surgery, University of Minnesota; Minneapolis, MN, USA; Schulze Diabetes Institute, University of Minnesota; Minneapolis, MN, USA; Department of Mechanical Engineering, University of Minnesota; Minneapolis, MN, USA; Department of Mechanical and Aerospace Engineering, University of South Florida; Tampa, FL, USA; Department of Physiology & Biomedical Engineering Mayo Clinic; Rochester, MN, USA; Mayo Clinic Graduate School of Biomedical Sciences, Mayo Clinic, Rochester, MN, USA; Wendy Novak Diabetes Institute, Norton Healthcare; Louisville, KY, USA; Department of Pediatrics, Pediatric Research Institute, University of Louisville; Louisville, KY, USA

## Abstract

Pancreatic islet transplantation can restore endogenous insulin production and offers a potential cure for diabetes, but its clinical impact has been limited by the inability to preserve large numbers of functional islets for timely use. Cryopreservation could provide an on-demand supply, yet conventional methods cause ice formation, cell injury, and loss of insulin secretion. We developed CryoMesh, a vitrification platform that combines a low-toxicity cryoprotectant with a thermally conductive, biocompatible mesh to enable ultra-rapid cooling and rewarming for ice-free cryopreservation. This approach supports long-term, clinical-scale preservation of both human pancreatic and stem cell–derived islets while maintaining viability, architecture, mitochondrial integrity, and glucose-responsive insulin secretion. Human islets preserved for up to one year restored normoglycemia in diabetic mice, with complete recovery of function and no increase in immunogenicity or loss of potency. The combination of high viability, recovery, and functional preservation at clinical scale has not been achieved previously. By decoupling islet isolation and manufacture from transplantation, CryoMesh enables extended quality, potency, and safety testing, cost-effective batch production, and global banking and distribution. These capabilities remove a major barrier to curative cell therapy for diabetes and establish a generalizable strategy for preserving complex multicellular therapeutics.

**One Sentence Summary:** A clinical scale vitrification platform enables long-term banking of human pancreatic and stem cell–derived islets for transplantation.

## INTRODUCTION

More than a century after the discovery of insulin (*1*), even advanced delivery systems such as continuous glucose monitors, insulin pumps, and closed-loop systems fail to maintain durable glycemic control (*2*). Islet transplantation offers a potential functional cure but remains constrained by islet scarcity, lifelong immunosuppression, and limited long-term graft survival (*3–5*).

The Edmonton Protocol established the feasibility of clinical islet isolation and transplantation (*6*). Yet, most patients require multiple transplants and ultimately return to insulin therapy (*5, 7–9*). Achieving euglycemia often requires large islet doses (>10,000 islet equivalents (IEQ)/kg or >1 million IEQ/patient), which exceeds what most single donor isolations can provide (*6, 9, 10*). This necessitates multiple donor infusions, increasing procedural and immunosuppressive risks as well as healthcare costs (*3, 11, 12*).

Pooling islets from multiple donors could help, but limited donor supply, rapid deterioration in culture (restricted to 3–7 days (*13, 14*)), and time-intensive quality testing limit the approach. Regulatory approval for islet transplantation has been hindered by the short shelf-life of fresh islets, which precludes comprehensive quality, safety, and efficacy assessments (*15*). Although recently approved for clinical use by a single manufacturer, concerns remain about the adequacy of testing and how islets should be regulated (*16*).

Human pluripotent stem cell–derived islets (SC-islets) offer a potentially limitless alternative to native pancreatic islets and have shown safety and efficacy in recent clinical transplant trials (*17, 18*). However, they are expensive to produce, prone to batch variability, and rapidly lose cellular identity in culture (*19–21*).

Cryopreservation of either islets or SC-islets would enable long-term banking and on-demand availability, allowing transplantation to occur at a time and place of choice while allowing thorough quality assessment and recipient preparation. At ultralow temperatures (<-150°C), biological activity ceases, making biopreservation theoretically indefinite (*22*). However, conventional slow-cooling cryopreservation methods, while effective for isolated single cells, fail for 3D multicellular constructs such as islets due to ice crystal formation (*23*).

Vitrification offers a promising alternative by enabling ice-free cryopreservation through rapid cooling into a *glass-like* state in the presence of cryoprotective agent (CPA) (*24, 25*). The challenge is balancing cooling and rewarming rates with CPA concentration: too little CPA or too slow cooling/rewarming leads to ice, while high CPA concentrations introduce toxicity. Thus, successful vitrification requires ultrafast cooling/rewarming, less toxic CPAs, or both. Further, due to the high concentrations of CPA solutions, loading and unloading protocols must be optimized to avoid osmotic stress. To address these trade-offs, numerous approaches have been explored, including Cryotop devices, hollow fibers, nylon meshes, droplet vitrification, silk scaffolds, cryotubes, and nanowarming (*23, 26–35*). Yet none has achieved the required combination of viability, recovery, function, and scale for clinical application.

Here, we introduce an engineered solution that enhances heat and mass transfer, enabling clinical-scale vitrification and rewarming (VR) of human pancreatic islets and human SC-islets. By mitigating ice formation and CPA toxicity, our approach offers a stable, on-demand supply of high-quality islets. For donor derived islets, this could enable elective scheduling, rigorous pre-transplant testing, immune tolerance protocols, global sharing/distribution, and better donor-recipient matching. For SC-islets, it further enhances manufacturing scalability, lowers costs, and ensures product consistency (Fig. 1). Together, these capabilities position cryopreservation via CryoMesh VR as a key enabler of scalable and durable cell-based therapies for diabetes.

**Fig. 1.**
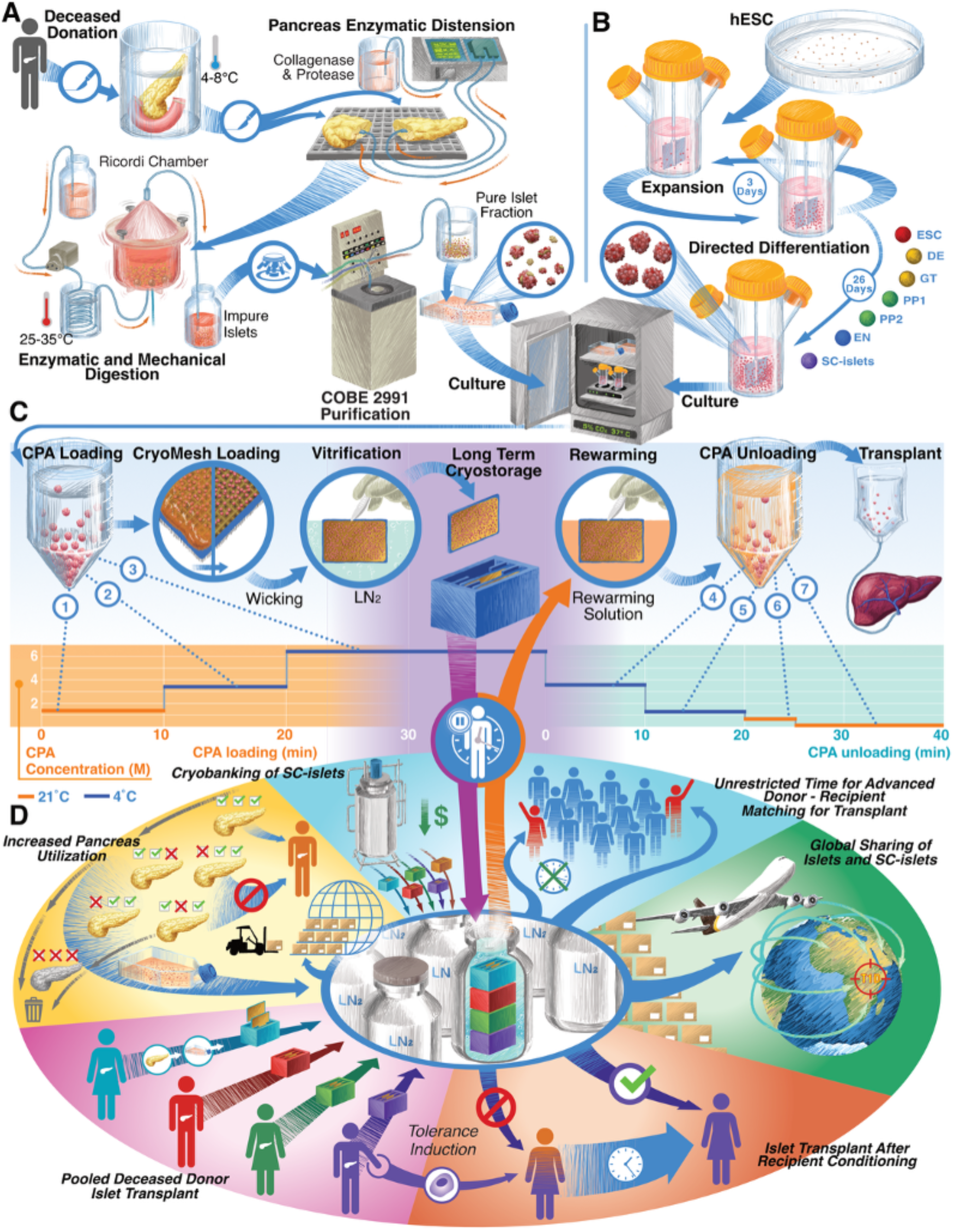
Process overview of pancreatic and stem cell-derived islet production, cryopreservation, and therapeutic benefits. (**A**) Human pancreatic islets were isolated from donor pancreases by enzymatic distension of the head and tail, digestion in a Ricordi chamber with agitation, and density-gradient purification (*36*). (**B**) Human embryonic stem cells were expanded and differentiated into stem cell–derived β cell islets (SC-islets) using a six-stage growth factor protocol (*19*). (**C**) Vitrification on gold-coated copper (Cu-Au) CryoMesh involved stepwise CPA loading, transfer of islets or SC-islets onto the mesh, removal of excess CPA by wicking, rapid plunging into liquid nitrogen, long-term cryobanking, and stepwise CPA unloading after rewarming. (**D**) Potential clinical benefits of islet cryopreservation include increased donor organ utilization, cryobanking of large islet or SC-islets batches as single-patient doses, improved donor–recipient matching, pooled donor transplantation, flexible scheduling of procedures, more time for batch quality assessment and safety testing, opportunities for immune tolerance induction, and global distribution. CPA, cryoprotective agent; hESC, human embryonic stem cell; LN_2_, liquid nitrogen

## RESULTS

For this study, we used both human native deceased donor pancreatic islets and human SC-islets. Islets were isolated from pancreata of brain-dead deceased donors deemed unsuitable for whole pancreas transplantation, using our GMP-compliant protocol involving enzymatic digestion and density gradient enrichment (Fig. 1A (*36*)). Isolation yields ranged from 554,489 to 737,322 IEQ (Table S1). Islet quality was confirmed by assessments of purity, sterility, endotoxin, viability, oxygen consumption rate (OCR), and glucose-stimulated insulin secretion (GSIS). High-purity fractions were used and 20–40% of the isolation as controls, with the remainder vitrified (351,033–585,361 IEQ; mean 452,936 ± 99,388).

SC-islets were generated by differentiation of Harvard University Embryonic Stem Cell Line-8 (HUES-8) pluripotent stem cells through a six-stage 26-day differentiation protocol (Fig. 1B). Differentiation was confirmed by flow cytometry to identify stage- and cell type–specific markers (Fig. S1). The fully differentiated insulin/ c-peptide positive SC-islet batches ranged from 189,601 to 478,223 IEQ (mean 245,456 ± 91,338), corresponding to 2.76×10^8^ to 7.17×10^8^ cells. Smaller batches were used for protocol development and validation.

### CryoMesh platform enables scalable vitrification of human islets and SC-islets

An overview of the CryoMesh VR process is shown in Fig. 1C. The CPA, composed of 22 wt% ethylene glycol (EG) and 22 wt% dimethyl sulfoxide (DMSO), was selected for its ability to prevent ice formation at the achieved cooling rates, with tolerable toxicity (*23*). To reduce osmotic stress and toxicity from the high CPA concentration, we used stepwise loading and unloading protocols (Fig. S2). A mass transport model with a toxicity cost function was employed to optimize the number, size, duration, and temperature of these steps (*23*). Microscopy confirmed ice-free vitrification: islets with sufficient CPA remained transparent, while those with insufficient CPA turned opaque due to ice formation (Fig. 2C).

**Fig. 2.**
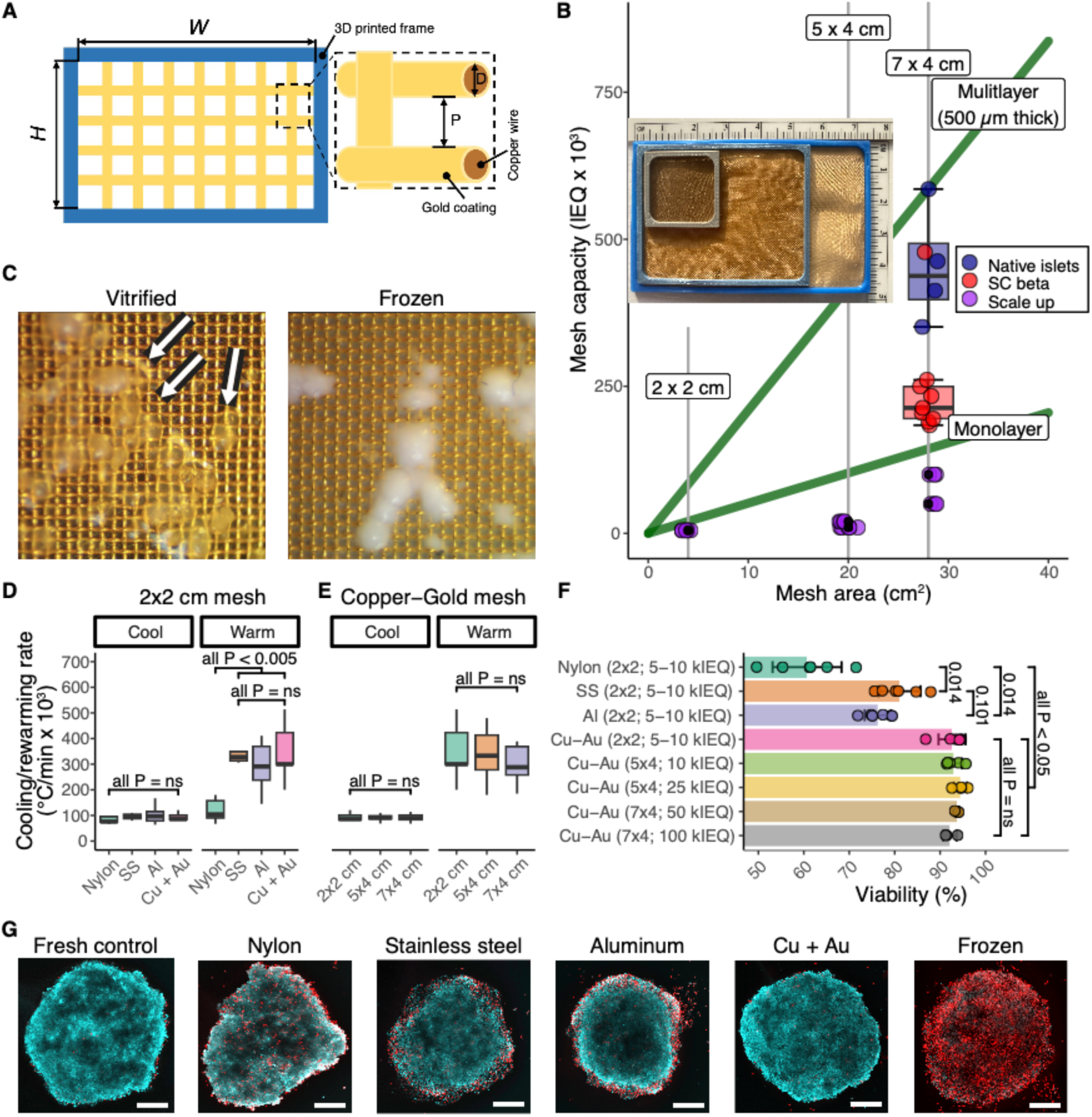
Engineering and performance of the CryoMesh system. (**A**) Schematic of the CryoMesh design. Wire diameter (D) is 50 ± 5 µm, and pore size (P) is 50 ± 5 µm. (**B**) Loading capacity of 2×2 cm, 5×4 cm, and 7×4 cm meshes. Green lines indicate theoretical monolayer capacity for 150 µm islets; multilayer capacity extends to 500 µm thickness. Purple points show SC-islet quantities tested during development; red (SC-islet) and blue (islet) points and boxplots indicate quantities tested on 7×4 cm meshes (inset photographs of mesh formats). (**C**) Magnified images of islets vitrified on Cu-Au CryoMesh (transparent, arrows, left) versus frozen (opaque, right). Mesh pore size = 50 µm. (**D**) Cooling and rewarming rates for mesh materials (n = 6–9 per group; Kruskal–Wallis with pairwise Wilcoxon tests). (**E**) Cooling and rewarming rates across mesh sizes (n = 9–14 per group; one-way ANOVA). (**F**) Post-rewarming viability of SC-islet on different mesh types (n = 3–6 per group; Kruskal–Wallis with pairwise Wilcoxon tests). (**G**) Representative viability (AO/PI) confocal images of rewarmed SC-islet (AO, live cells in cyan; PI, dead cells in red). Scale bar, 75 µm. Data shown as box-and-whisker plots or mean ± SD. Al, aluminum; AO, acridine orange; Cu + Au, copper-gold; IEQ, islet equivalents; ns, not significant; PI, propidium iodide; SC, stem cell; SS, stainless steel.

After CPA loading, islets or SC-islets were placed on a mesh support, and excess CPA was wicked off to reduce thermal mass. The mesh was then rapidly plunged into liquid nitrogen (LN₂) for vitrification (Fig. 1C) and stored at cryogenic temperatures. For rewarming, meshes were removed from LN₂, immersed in rewarming solution, and subjected to stepwise CPA unloading. The samples were then cultured for 3–16 hours before use.

### High-conductivity metal mesh facilitates rapid and uniform heat transfer

In preliminary studies, scaling nylon CryoMesh from 2,500 to ≥10,000 IEQ reduced islet viability and recovery (Fig. 2F-G, (*23*)), likely due to uneven cooling and limited convective heat transfer efficiency, which promoted ice formation (*23, 35*). To address these issues, we made two key changes: 1) plunging the mesh vertically during cooling decreased the insulating vapor layer (Leidenfrost effect) that typically forms around objects submerged in liquid nitrogen, thereby accelerating cooling rates, and 2) switching from nylon to engineered metal meshes improved heat transfer by combining conduction with convective heat transfer. These modifications kept temperature variation below 6% across a 5×4 cm CryoMesh (*35*).

To further characterize the heat transfer, we compared 2×2 cm meshes of nylon, stainless steel, aluminum, and gold-coated copper (Cu–Au) using <500-µm-thick, CPA-loaded SC-islet layers containing 5,000–10,000 IEQ (Fig. 2D). The Cu–Au mesh cooled at 9.5×10⁴ °C min⁻¹, 17% faster than nylon, and showed lower variability than aluminum (coefficient of variation, CV, 91% higher) and stainless steel (CV 36% higher). During rewarming, Cu–Au reached 3.48×10⁵ °C min⁻¹, 2.9× faster than nylon, 18% faster than aluminum, and within 2% of stainless steel. Because the stainless-steel wires were thinner (30 µm vs 50 µm for Cu–Au and aluminum), reducing Cu–Au wire diameter may further improve performance by enhancing heat transfer (*35*).

Cu-Au provided the highest post-thaw viability across multiple mesh sizes, reaching up to 96.1% (mean 92.6 ± 3.0%; Fig. 2F), surpassing stainless steel (81.1 ± 4.6%), aluminum (76.3 ± 2.9%), and nylon (60.7 ± 7.6%). Live–dead staining showed that Cu-Au-preserved islets closely resembled fresh controls, while nylon resulted in dispersed cell death (suggesting ice formation). Other metals showed signs of peripheral injury (indicating toxicity from surface interaction) (Fig. 2G). Due to its superior thermal uniformity, high viability, scalability, and gold-mediated biocompatibility, we selected the Cu-Au CryoMesh as the platform for clinical-scale islet and SC-islet vitrification. This enabled up to a 10,000-fold increase in throughput compared to alternatives like laser “nanowarming,” which achieved rewarming rates of up to 10 M °C/min in µL sized droplets with 10’s of islets in each (Table S2, (*23, 35*)).

### Large-format CryoMesh preserves clinical-scale islet and SC-islet batches

We set a target capacity of 250,000–500,000 IEQ per CryoMesh, potentially combining multiple meshes for clinical-scale batches. To optimize size, we tested 2×2 cm, 5×4 cm, and 7×4 cm meshes (Fig. 2A and 2B inset) and successfully vitrified SC-islet test batches of 5,000–100,000 IEQ (Fig. 2B, purple points) without visible ice formation.

To estimate maximum loading, we modeled the optimal packing of 150 µm spheres (size of an IEQ) (*37*). Monolayer packing supported 5,133 IEQ/cm², or ∼143,000 IEQ on a 7×4 cm mesh (Fig. 2B, green line; Fig. S4). Multilayer packing up to 500 µm thickness, beyond which ice formation occurred (Fig. S5), increased capacity to 20,925 IEQ/cm², or ∼585,900 IEQ per 7×4 cm mesh. The natural size variability of islets and SC-islets (Fig. S3) may further increase this capacity due to more efficient packing.

After establishing a VR protocol that preserved high post-thaw viability in SC-islets, we tested the Cu-Au CryoMesh at clinically relevant scales. Using 7×4 cm meshes, we vitrified and rewarmed up to 585,361 IEQ (8.78×10⁸ cells) of native human islets and 478,223 IEQ (7.17×10⁸ cells) of SC-islets on single CryoMeshes (Fig. 3B), consistent with the theoretical capacity of ∼600,000 IEQ (∼9 X 10^8^ cells) per mesh (Fig. 2B, figs. S4–S5). This capacity (∼600,000 IEQ) exceeds most clinical islet isolations and corresponds to a therapeutic dose for adult recipients.

**Fig. 3.**
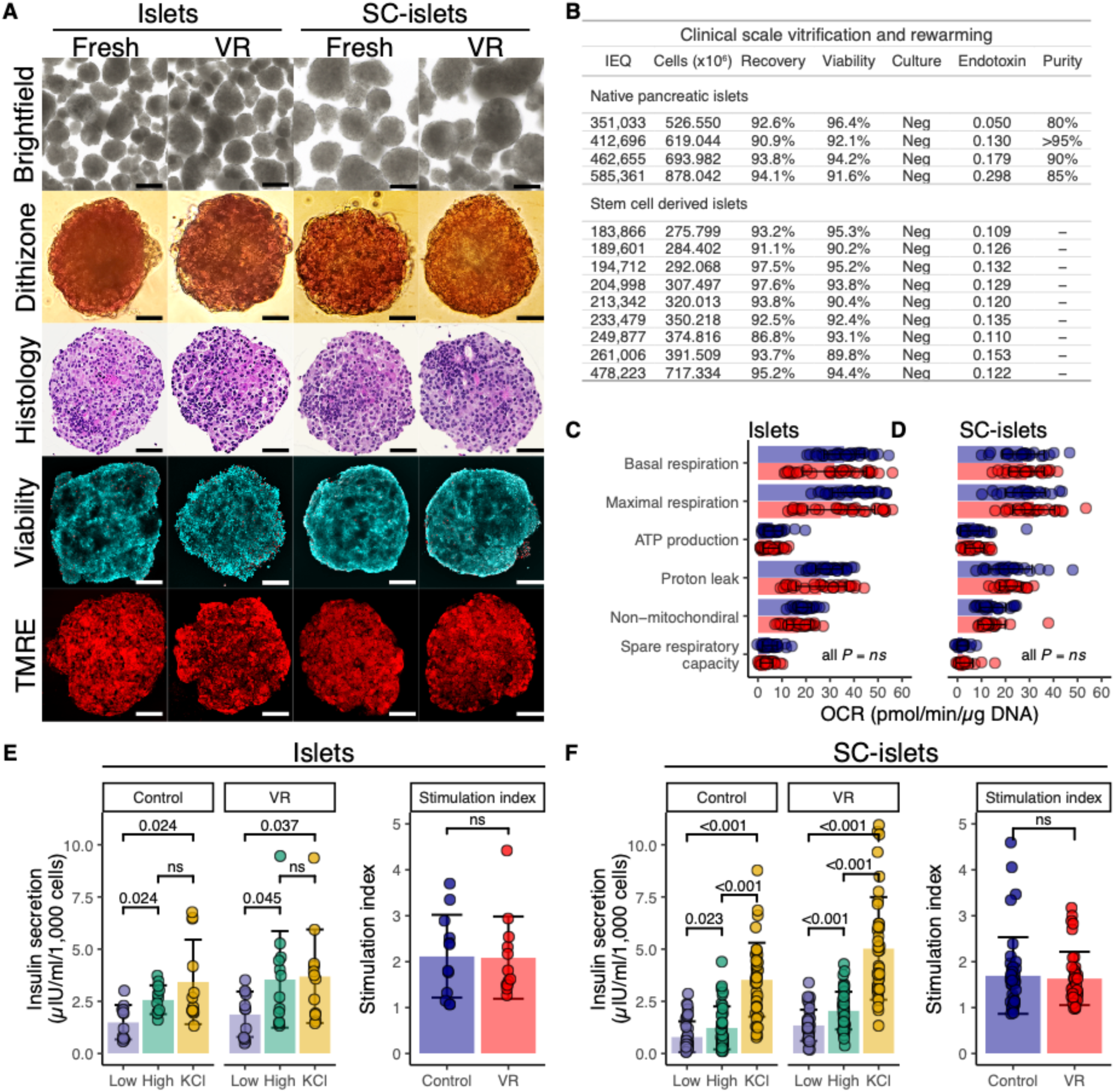
Biological assessment of vitrified and rewarmed pancreatic islets and SC-islets. (**A**) Brightfield microscopy, dithizone staining of β-cell zinc granules, hematoxylin and eosin histology, AO/PI viability staining (AO, live cells in cyan; PI, dead cells in red), and TMRE analysis of mitochondrial membrane potential of fresh or vitrified and rewarmed (VR) islets and SC-islets. Scale bars: 100 µm for brightfield; 25 µm for Dithizone and Histology; 75 µm for Viability and TMRE. (**B**) Clinical-scale VR outcomes include total IEQ and cell quantities processed; recovery, viability, and sterility (5-day bacterial culture; peak endotoxin levels (EU/mL) measured at 0-, 3-, and 24-hours post-rewarming); and purity of pancreatic islets. (**C, D**) Mitochondrial respiration in fresh versus VR samples was measured using Seahorse Mito Stress Test. Oxygen consumption rates and functional metrics for (**C**) native islets and (**D**) SC-islets; n = 24–37 per group. Statistical tests: Kruskal–Wallis with pairwise Wilcoxon comparisons. (**E–F**) *In vitro* glucose-stimulated insulin secretion (GSIS) responses and stimulation indices for fresh and VR islets (**E**) and SC-islet (**F**). Data are presented as box-and-whisker plots or mean ± SD. AO, acridine orange; EU, endotoxin units; IU, international units; ns, not significant; OCR, oxygen consumption rate; PI, propidium iodide; SC, stem cell; TMRE, tetramethylrhodamine ethyl ester; VR, vitrified and rewarmed.

### Vitrification preserves islet and SC-islet morphology, viability, mitochondrial health, and glucose-responsive insulin secretion in vitro

Post-VR morphology was indistinguishable from fresh controls, as shown by brightfield imaging, dithizone staining of zinc-rich β cells, and histology (Fig. 3A). No increase in intra- or intercellular white space was detected, consistent with the absence of ice formation (*38*). Live-dead staining confirmed high overall viability (Fig. 3A, Fig. S6), while tetramethylrhodamine ethyl ester (TMRE) staining demonstrated preserved mitochondrial membrane potential in SC-islets and a slight reduction in native islets at 3 h post-VR (Fig. 3A, figs. S7-S8).

Quantitative outcomes demonstrated 93.6 ± 2.2% viability for native islets and 92.7 ± 2.2% for SC-islets after VR (Fig. 3B). Recovery rates were also high (92.9 ± 1.4% and 93.5 ± 3.3%, respectively). All samples remained sterile, and endotoxin levels measured repeatedly over 24 hours post-rewarming were below the clinical threshold for islet release (<5 EU/mL) (*39*). Seahorse Mito Stress Tests confirmed no significant differences in oxygen consumption rate (a key predictor of posttransplant function (*39*)) or other mitochondrial parameters, including basal respiration, ATP production, maximal respiration, and spare capacity, between fresh and VR-treated samples (Figs. 3C–D, Fig. S9).

Functional recovery after VR was assessed *in vitro* using a glucose-stimulated insulin secretion (GSIS) assay. VR-treated islets and SC-islets responded to high glucose and KCl depolarization, with stimulation indices comparable to fresh controls (Figs. 3E–F), confirming preserved glucose responsiveness.

### Cryopreserved islets and SC-islets restore durable glucose control in vivo

We next evaluated *in vivo* function using a xenotransplantation model in non-diabetic immunodeficient SCID-beige mice. Human islets or SC-islets were transplanted under the kidney capsule of recipient mice, and human insulin levels were measured in recipient serum eight weeks later. Both fresh and VR-treated grafts secreted basal circulating human insulin, which increased after intraperitoneal glucose administration, confirming preserved glucose responsiveness (Figs. 4A–D). Confocal imaging of explanted grafts showed strong insulin and glucagon staining in both groups, indicating intact β- and α-cells (Figs. 4E–F; Fig. S10).

**Fig. 4.**
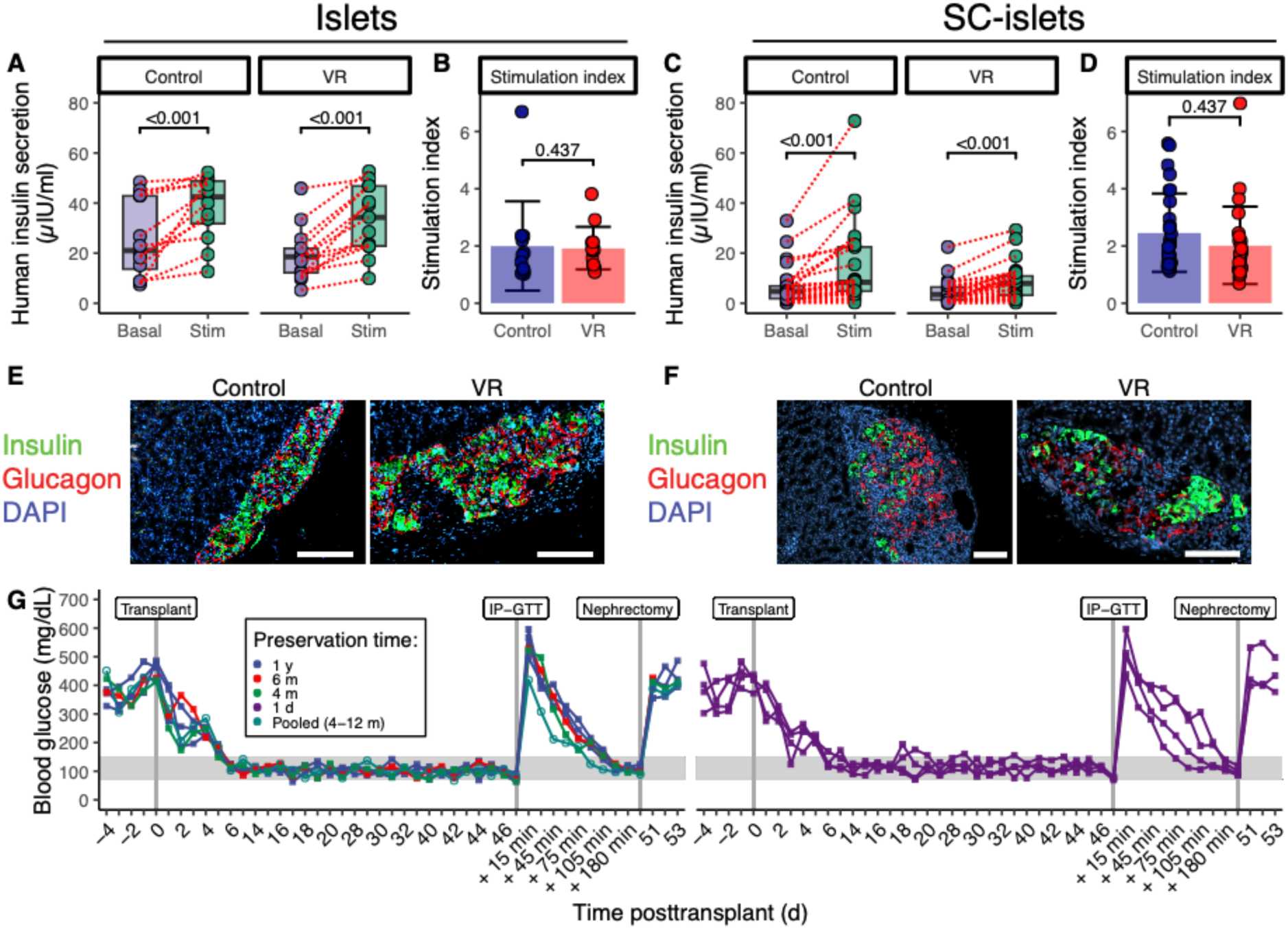
In vivo function of vitrified and rewarmed islets and SC-islets. In all panels, native pancreatic islets (left) and SC-islets (right) are shown. (**A–D**) Human islet and **SC-islet** xenotransplantation in immunodeficient (SCID beige) mice: basal and intraperitoneal (IP) glucose-stimulated human serum insulin levels and corresponding stimulation indices. (n = 12–28 per group; paired Wilcoxon test). (**E-F**) Confocal images of explanted grafts stained for insulin (red), glucagon (green), and nuclei (DAPI, blue). Scale bars = 100 µm. (**G**) Reversal of hyperglycemia in chemically diabetic immunodeficient (NSG) mice transplanted with cryopreserved human islets (left) or SC-islets (right) stored for 1 day to 1 year. An intraperitoneal glucose tolerance test (IP-GTT) was performed on day 49, followed by nephrectomy of the graft-bearing kidney on day 50. Gray shading indicates the normoglycemic range. n = 5 for islet transplant and n = 4 for SC-islet grafts. Data are shown as box-and-whisker plots or mean ± SD. Abbreviations: DAPI, 4′,6-diamidino-2-phenylindole; NSG, NOD scid gamma; SC, stem cell; VR, vitrified and rewarmed.

Therapeutic efficacy was then assessed in chemically (streptozotocin) induced diabetic immunodeficient NSG mice transplanted with VR-preserved human islets or SC-islets stored from 1 day to 1 year (Fig. 4G), including a pooled four-donor islet preparation. All recipients achieved normoglycemia within 6 days and maintained tight glycemic control for 50 days. Glucose tolerance tests on posttransplant day 49 showed transient hyperglycemia, with basal levels restored within 180 minutes. Nephrectomy of the graft-bearing kidneys led to rapid recurrence of hyperglycemia, confirming graft-dependent euglycemia.

### Vitrification induces minimal transcriptional change and stress response

We examined whether VR altered RNA expression or cellular pathways by performing bulk RNA sequencing on four different human islet samples (from various donors) and four SC-islet batches, both before and after VR. Unsupervised clustering revealed that native islets grouped by donor rather than treatment, with no differences observed between fresh and VR samples (Fig. S10A). In contrast, SC-islets results clustered according to treatment group (fresh vs. VR) rather than differentiation batch (Fig. S10B), a pattern confirmed by multidimensional scaling (Fig. S11).

Differential gene expression (DGE) analysis revealed no changes between fresh and VR-treated islets (absolute log₂ fold change >1, p < 0.05, FDR <7.5%; Fig. 5A, Table S3). In SC-islets, 20 transcripts were significantly upregulated after VR (Figs. 5B–C; Table S4), enriched in pathways related to cellular stress, apoptosis, and NF-κB regulation (Table S5). Pathway analyses supported these findings: gene ontology (GO) terms were enriched for apoptosis, cytokine signaling, and stress responses (Fig. 5D), while gene set enrichment analysis (GSEA) confirmed activation of TNFα–NF-κB signaling, inflammation, hypoxia, and apoptosis (Fig. S12). Further description of specific gene alteration is presented in the Supplemental Information.

**Fig. 5.**
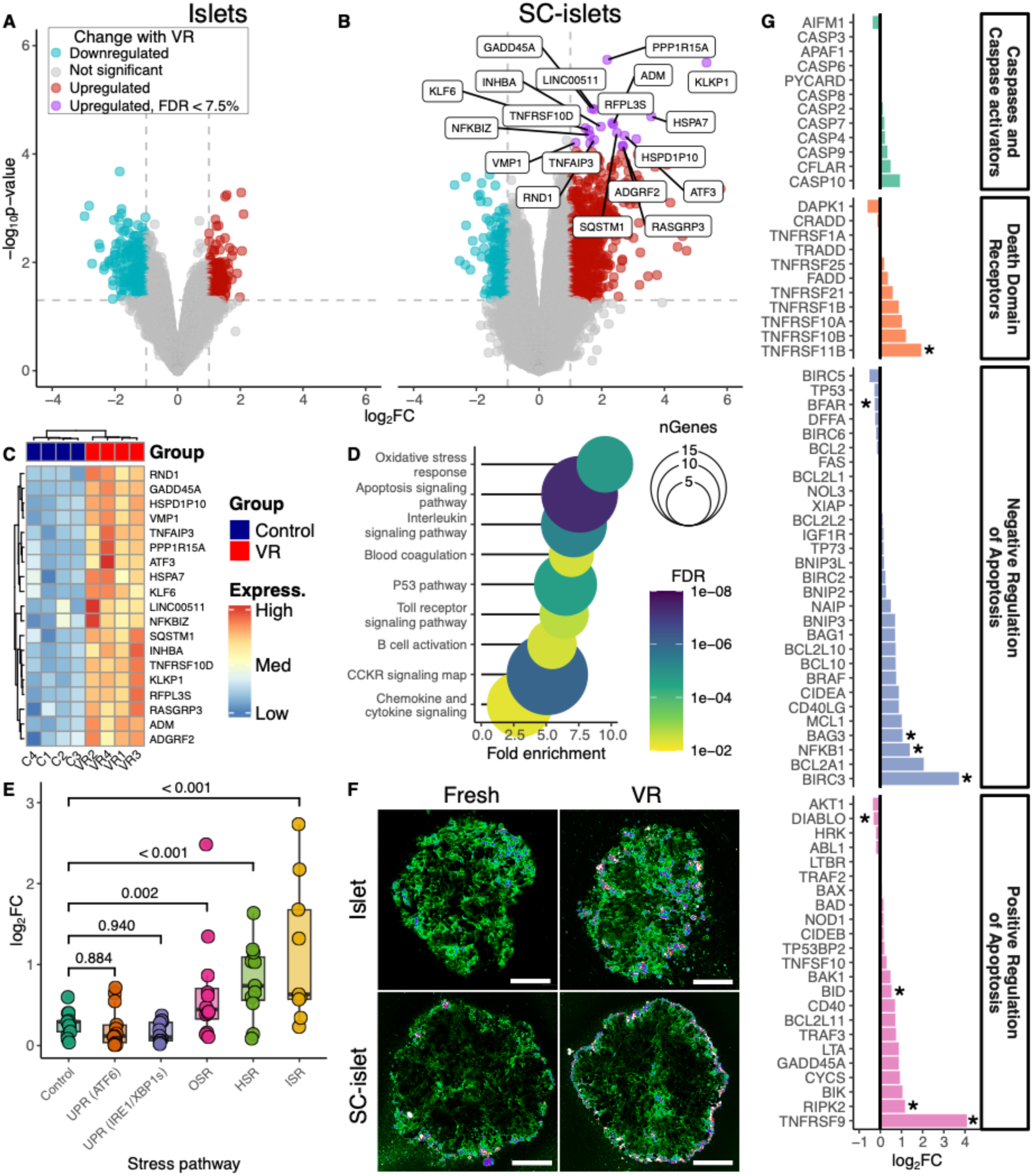
Transcriptomic and stress responses to vitrification and rewarming. (**A, B**) Bulk RNA-seq volcano plots comparing fresh versus VR native islets (**A**) and SC-islets (**B**). (**C**) Heatmap of differentially expressed genes in VR versus control SC-islets. (**D, E**) Pathway analyses highlighting enrichment of apoptosis, cytokine signaling, and stress responses in SC-islets (PANTHER categories; box-and-whisker plots). (**F**) Confocal images of Annexin V staining in VR islets and SC-islets, indicating apoptosis. Scale bar = 75 µm. (**G**) qPCR analysis of apoptosis-related gene expression in VR-treated SC-islets (mean log₂ fold change [log₂FC], n = 3–4 per group; Student’s *t*-test, * = p < 0.05). Abbreviations: FDR, false discovery rate; HSR, heat shock response; ISR, integrated stress response; OSR, oxidative stress response; SC, stem cell; UPR, unfolded protein response; VR, vitrified and rewarmed.

To assess stress at the cellular level, we measured canonical pathways. VR upregulated oxidative, heat shock, and integrated stress responses (Fig. 5E), while unfolded protein response genes were unchanged. Confocal imaging showed modest increases in membrane-bound Annexin V after VR in both islets and SC-islets, indicating a small apoptotic cell population (Fig. 5F). qPCR analysis further revealed that VR upregulated both pro- and anti-apoptotic genes in SC-islets (Fig. 5G). No significant transcriptional changes were observed in islets (all p > 0.05).

### Cryopreservation preserves native immunological profiles of islets and SC-islets

Instant blood-mediated inflammatory response (IBMIR) is a rapid innate immune reaction triggered when transplanted islets come into contact with blood and can destroy up to ∼60% of grafted tissue (*40, 41*). To determine whether vitrification alters antigenicity or immunogenicity, we performed an *in vitro* IBMIR assay comparing fresh and vitrified–rewarmed (VR) pancreatic islets and stem cell–derived islet (SC-islet) clusters. Whole blood from donors with type 1 diabetes (not receiving immunosuppression) was incubated with islets or SC-islets and analyzed using 25-marker spectral flow cytometry (Fig. S13), with blood-only samples serving as controls. Major innate immune populations, including neutrophils, monocytes, dendritic cells (DCs), myeloid-derived suppressor cells (MDSCs), NK cells, and NKT cells, were identified by conventional gating (Fig. S14), followed by UMAP-based dimensionality reduction and AI-assisted phenotypic annotation.

Analysis of the neutrophil compartment revealed a functional continuum spanning twelve distinct phenotypic clusters (Fig. 6A-C). These ranged from quiescent populations (CD62L^high^, CD11b^low^) to primed and fully activated effector cells, characterized by degranulation (CD66b^high^), activation (CD62L^low^) and hyper-adhesion (CD54^high^) (Fig. 6D, Table S6). As expected, exposure to fresh human islets triggered a shift from the quiescent state toward activated phenotypes compared to no-islet controls (Fig. 6B, E). Importantly, vitrification and nanowarming did not exacerbate this inflammatory response (Fig. 6B, E, G). SC-islets exhibited a comparable safety profile. While SC-islets induced a modest upregulation of specific hyper-adhesive neutrophil subsets (Cluster 10), vitrification did not amplify this response (Fig. 6C, F, H).

**Fig. 6.**
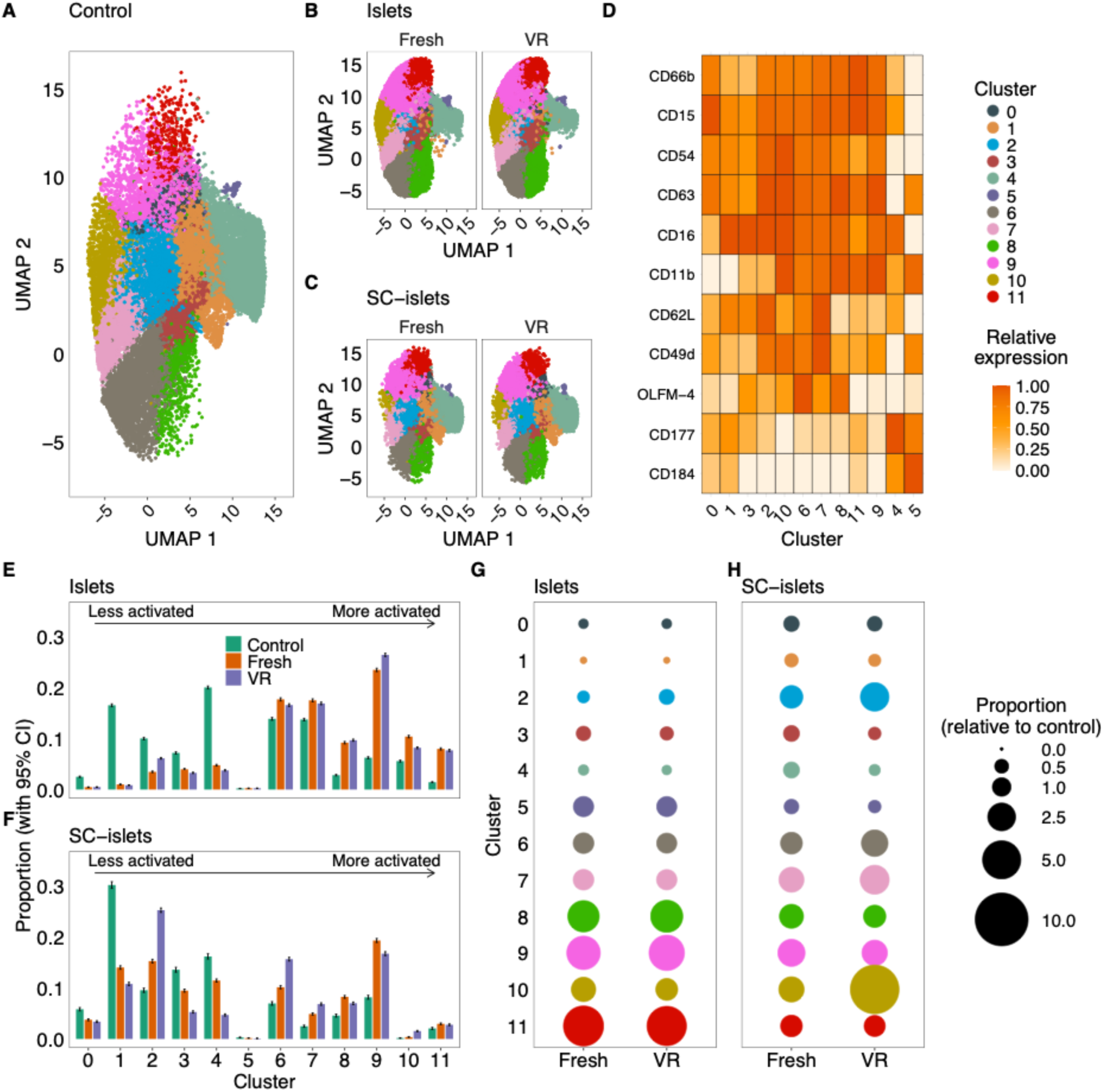
Instant blood-mediated immune responses (IBMIR) in neutrophils. Dimensional reduction analysis depicting cell subpopulation clusters for (**A**) control blood samples, (**B**) fresh and vitrified/rewarmed (VR) islets, and (**C**) fresh and VR SC-islets. (**D**) Transformed and scaled relative expression of key cell surface markers identified in **A**-**C**. Population sizes for control blood samples, fresh islets/ SC-islets, and VR islets/SC-islets are shown for each cluster in (**E**) islets and (**F**) SC-islets. Clusters are ordered from lowest activation state (cluster 0) to highest (cluster 11). Data are mean ± 95% confidence interval. Bubble plots illustrate the relative population size compared to control for (**G**) fresh and VR islets and (**H**) fresh and VR SC-islets.

Parallel high-dimensional analysis of the mononuclear phagocyte compartment resolved seven distinct populations (Clusters 0–6) (Fig. S15, Table S7). The monocyte lineage displayed a clear activation gradient, ranging from patrolling non-classical monocytes (Cluster 0) to inflammatory classical monocytes (Cluster 2). Notably, we identified a highly activated classical subset (Cluster 3), distinguished by maximal expression of the degranulation marker CD63, and a transitioning intermediate subset (Cluster 1) enriched for CD177. Distinct from the monocyte lineage, we observed quiescent pDCs (Cluster 4), mature antigen-presenting pDCs (Cluster 5; HLA-DR^high^), and a population of granulocytic-MDSCs (Cluster 6) characterized by a hyper-adhesive (CD54^high^), suppressive phenotype (HLA-DR^neg^).

Exposure of blood to islets led to some remodeling of monocyte phenotypes, characterized by a relative decrease in inflammatory classical monocytes (Cluster 2) and concomitant increases in transitioning intermediate monocytes (Cluster 1) and activated classical monocytes (Cluster 3) (Fig. S15, Table S7). In contrast, NK and NKT cell populations remained stable across all conditions for islets (Figs. S16-S17, Tables S8-S9). In SC-islets, there was a minor shift from resting NKT cells (Cluster 1) to activated effectors (Cluster 2) (Fig. S17F, Table S9). For all identified populations, including the activated neutrophil effectors and remodeled monocyte subsets, no notable differences were observed between fresh and VR islets.

Adaptive immunity was tested in a murine allogeneic transplant model (Fig. 7). Fresh islets were rejected after an average of 12 days, while VR-preserved islets showed a modest, non-significant extension to 14 days.

**Fig. 7.**
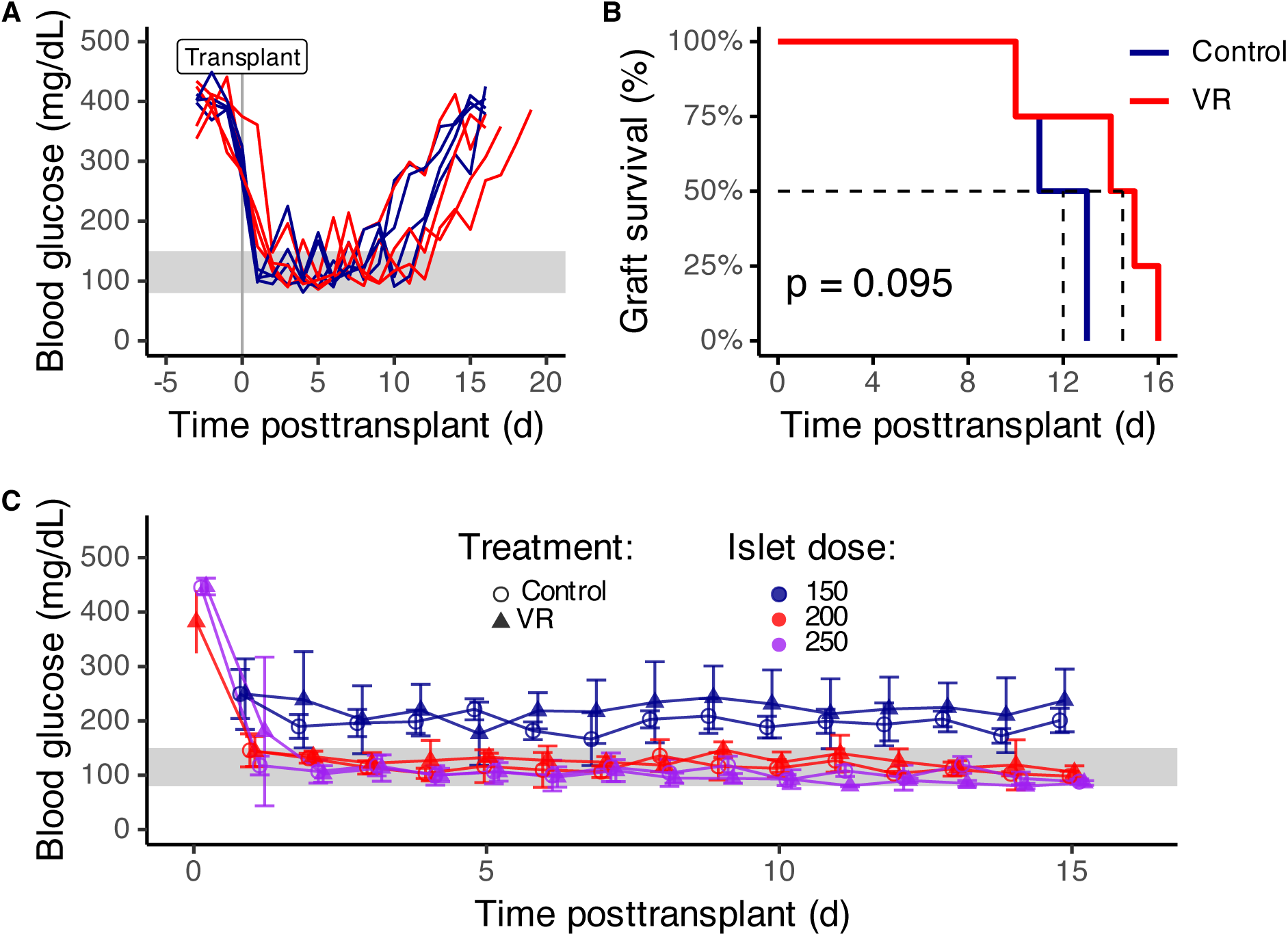
Alloimmune rejection and functional bioequivalence of mouse islets in a streptozotocin-induced diabetes transplant model. (**A**) Blood glucose levels in islet transplant recipients measured from 3 days before to 20 days after transplantation. The gray shaded region denotes the normoglycemic range. (**B**) Kaplan–Meier analysis of graft survival, with median survival times indicated by dashed lines. Control (blue) and vitrified/rewarmed (VR; red) recipients were compared using a log-rank test (n = 4 per group). (**C**) Limiting-dose syngeneic islet transplants were performed in streptozotocin-induced diabetic C57BL/6 mice to determine the minimal islet mass required to restore normoglycemia. Fresh and VR islets reversed diabetes at comparable doses, demonstrating functional bioequivalence. The gray shaded region indicates the target normoglycemic blood glucose range.

### Vitrified islets are functionally bioequivalent to fresh grafts

To evaluate the biological potency of vitrified and rewarmed (VR) islets, we performed syngeneic transplants in C57BL/6 mice. Consistent with previous findings (*23*), 250 VR islets reversed diabetes. Both fresh and VR islets restored normoglycemia at 200 islets per mouse, but not at 150, establishing a shared minimum effective dose between 150 and 200 islets (Fig. S21). These results confirm the *in vivo* efficacy and functional bioequivalence of VR and fresh islets.

## DISCUSSION

Islet transplantation is at a pivotal point in diabetes treatment. Long-term studies have shown sustained improvements in quality of life, near elimination of severe hypoglycemic events, improved efficacy compared with insulin, and reduced long-term healthcare costs (*4, 7, 8, 42*). However, securing sufficient islet mass for lasting insulin independence remains a central challenge, leading to strategies such as pooled donor transplants and SC-islet cell therapies (*19*). A decade after their development, SC-islets have restored normoglycemia in clinical trials (*17, 18*), but both native and stem cell–derived sources are constrained by short preservation windows. To overcome these challenges, we developed a clinical-scale vitrification platform for long-term cryopreservation of islets and SC-islets.

Using a Cu-Au CryoMesh with a low-toxicity EG/DMSO cryoprotectant, we achieved ultra-rapid cooling and rewarming rates sufficient to prevent ice formation in large batches (>500,000 IEQ per mesh). VR islets remained viable for at least one year in storage, maintaining high recovery, preserving insulin secretion and mitochondrial function, and exhibiting minimal transcriptomic and immunogenic changes. *In vivo*, both islets and SC-islets subjected to VR reversed diabetes after transplantation, confirming preserved therapeutic efficacy. Across more than 4 million IEQ, all preparations met clinical release criteria for viability, purity, identity, potency, and sterility.

Existing cryopreservation methods have not achieved the ideal combination of viability, recovery, function, and clinical scalability. Slow-freezing suffers from intracellular ice formation, CPA toxicity, and thermal stress, leading to unacceptable post-thaw results (*43*). Large-scale freezing preserved tens of thousands of islets but with limited functionality (*44, 45*). Other techniques improved viability but relied on animal-derived reagents or nonclinical processes (*46–49*). Vitrification attempts by others failed to reach sufficient thermal rates to avoid ice formation, especially during warming, or required toxic CPA levels, restricting their use to small volumes with incomplete recovery (*50–54*). Alternative devices like droplet systems, hollow fibers, nylon meshes, silk scaffolds, and nanowarming have made incremental advances but still lack the reliability and scalability needed for clinical application (*23, 28–32, 34*).

The Cu-Au CryoMesh addresses these challenges by combining high thermal conductivity, biocompatibility, and manufacturing simplicity. It enables long-term storage without toxicity, achieves ultrarapid cooling and rewarming rates, and supports over 500,000 IEQ (∼0.8×10^9^ cells) per 7×4 cm mesh, with the option to scale further using multiple or larger meshes. By preserving both islets and SC-islets for at least one year, the platform makes high-dose, pooled-donor, single-transplant procedures possible, reducing risks associated with marginal doses, multiple procedures, and repeated induction immunosuppression. It also offers flexible scheduling, extended quality and safety testing, and standardized batch validation—especially valuable for SC-islets manufacturing. Additionally, long-term banking enables improved donor–recipient matching and fosters national and international sharing, ultimately lowering costs, increasing pancreas utilization, and enhancing patient access and transplant outcomes. Since it does not alter the immunological landscape, VR islets and SC-islets provide a platform for recipient conditioning and tolerance induction, supporting immunosuppression-free islet transplantation.

Study limitations include reliance on direct LN2 contact, which could be mitigated with sterile LN2 or a sealed system; a non-hermetic storage format, although no contamination was observed; and limited biochemical validation of transcriptomic data. Although the long-term effects of trace Cu-Au exposure at ultralow temperatures in a vitrified state did not show toxicity, further evaluation for biocompatibility and FDA compliance is necessary.

In summary, the Cu-Au CryoMesh establishes a clinically compatible, clinical-scale platform for cryopreserving islets and SC-islets, while maintaining function and safety for at least one year. This advancement enables elective, high-dose transplantation, better donor–recipient matching, and global sharing of donor material. It also facilitates standardized manufacturing and batch validation of SC-islets. By providing long-term, on-demand access to transplant-ready islets, this technology removes a significant barrier to the widespread use of beta cell replacement therapy, bringing the field closer to a lasting cure for diabetes.

## MATERIALS AND METHODS

### Human pancreas recovery, islet isolation, and islet culture

Human pancreata from deceased donors who had consented to research but were declined for clinical transplantation were recovered by LifeSource organ procurement organization (OPO, Minneapolis, MN) from within its service area (Minnesota, North Dakota, South Dakota, and parts of western Wisconsin). Organs were flushed in situ with University of Wisconsin (UW) solution and shipped as pancreatico-duodenal grafts to the University of Minnesota. GMP-grade islet isolation was performed at the University of Minnesota Molecular and Cellular Therapeutics facility using described protocols (*36*), summarized below.

After surgical trimming, pancreata were perfused via the pancreatic duct with collagenase (CIzyme CI:CII, VitaCyte, Cat. #001-1000) and neutral protease (NB GMP Grade, SERVA/Nordmark, Cat. #30303). Digestion was performed in a Ricordi chamber, and islets were purified using continuous Ficoll density gradients on a cooled Cobe 2991 cell separator. Islets were washed and cultured in CMRL medium supplemented with 25% human serum albumin, 1000 U/mL heparin, and 10 μg/mL ciprofloxacin. They were plated at 25,000–27,500 IEQ per T-175 flask and incubated at 37°C with 5% CO₂ for 24 hours, then maintained at 22°C until cryopreservation.

### SC-islet directed differentiation from HUES-8 and flow cytometry-based characterization

Human embryonic stem cells (hESCs) from the HUES-8 line (Harvard University) were expanded and differentiated in suspension culture using stage-specific media (S1, S2, S3, and BE5) to generate SC-islets (*55*). HUES-8 cells were cultured in 500 mL spinner flasks (Corning, 3153) by seeding 150 million cells (5 × 10⁵ cells/mL) in mTeSR1 media (STEMCELL Technologies, 85850) with 10 μM Y27632 (R&D Systems, 1254) and maintained at 70 rpm in a humidified incubator at 37°C, 5% CO₂. Media were replaced after 48 hours with fresh mTeSR1 without Y27632. Cells were passaged every 72 hours by dissociation to single cells using Accutase (Millipore Sigma, A6964) and mechanical disruption, then resuspended in fresh mTeSR1 with Y27632 (*56*).

SC-islet differentiations were initiated three days after SC spheroid formation and expansion in mTeSR1 media inside a spinner flask. Clusters were allowed to settle by gravity, and the media was replaced with protocol-specific media containing appropriate growth factors. Cell differentiation was sequentially directed toward definitive endoderm, primitive gut tube, pancreatic progenitor 1, pancreatic progenitor 2, endocrine progenitors, and ultimately into SC-islets.

#### Directed differentiation

Differentiation of pluripotent stem cells into SC-islets was performed by changing media in the spinner flask and supplementing with small molecules and growth factors specific to each differentiation stage. Differentiation media forumulation and inductive agents are detailed in Supplementary Information. The media changes were as follows: *Day 1:* S1 media with 100 ng/mL Activin A plus 3 µM CHIR99021; *Day 2:* S1 media with 100 ng/mL Activin A; *Day 4:* S2 media with 50 ng/mL KGF; *Day 6:* S3 media with 50 ng/mL KGF, 250 nM Sant-1, 500 nM PDBu, 200 nM LDN 193189, 2 µM RA, and 10 µM Y27632; *Day 7:* S3 media with 50 ng/mL KGF, 250 nM Sant-1, 500 nM PDBu, 2 µM RA, and 10 µM Y27632; *Days 8, 10, and 12:* S3 media with 50 ng/mL KGF, 250 nM Sant-1, 100 nM RA, 10 µM Y27632, and 5 ng/mL Activin A; *Days 13 and 15:* BE5 media with 250 nM Sant-1, 20 ng/mL betacellulin, 1 µM XXI, 10 µM ALK5i, 1 µM T3, and 100 nM RA; *Days 17 and 19:* 20 ng/mL betacellulin, 1 µM XXI, 10 µM ALK5i, 1 µM T3, and 25 nM RA; *Days 20–26:* S3 media only *Characterization of SC-islets by flow cytometry:* Spheroids were dispersed into single cells by incubation in TrypLE Express at 37°C for 10 minutes and mechanically disrupted. Cells were quenched with S3 media, washed with PBS, fixed in 4% PFA for 60 minutes, and stored at 4°C in PBS until further use. For beta-cell-specific viability experiments, dissociated clusters were stained with live/dead dye (LIVE/DEAD Fixable Near-IR, ThermoFisher, Waltham, MA), washed, and fixed before staining. Prior to staining, cells were filtered through a 40 µm cell strainer, permeabilized in blocking buffer (1x PBS, 0.1% Triton X-100, 5% donkey serum) at room temperature for 40 minutes, and washed three times in PBST (0.1% Triton X-100). Cells were then incubated with primary antibodies in the blocking buffer for 1 hour at room temperature, washed three times with PBST, and incubated with secondary antibodies in blocking buffer for 1 hour at room temperature. After labeling, cells were washed three times, resuspended in PBST at a concentration of 1 × 10^6^ cells/mL, and analyzed using an Attune flow cytometer (ThermoFisher). Data were acquired and analyzed with NXT v. 3.1 (ThermoFisher) and FlowJo v.11 (FlowJo, Ashland, OR) software.

The following primary antibodies were used for flow cytometry: goat anti-PDX1 (R&D Systems, AF2419), mouse anti-NKX6.1 (DSHB, F55A12), rabbit anti-chromogranin (Abcam, ab15160), rat anti-c-peptide (DSHB, GN-ID4), and mouse anti-glucagon (Sigma, G2654). Secondary antibodies for flow cytometry included donkey anti-goat Alexa Fluor 488 (Invitrogen, A-11055, 1:1,000), donkey anti-rabbit Alexa Fluor 488 (Invitrogen, A-21206, 1:1,000), donkey anti-rat Alexa Fluor 488 (Invitrogen, A-21208, 1:1,000), and donkey anti-mouse Alexa Fluor 647 (Invitrogen, A-31571, 1:1,000).

### Quality control (IEQ estimation, endotoxin and culture-sensitivity tests)

Islet equivalence (IEQ), endotoxin levels, and culture sensitivity were measured for islets and SC-islets after islet isolation, SC-islet product release, before CPA loading (pre-vitrification), and immediately after CPA unloading (post-rewarming). Additionally, endotoxin levels in culture media were evaluated 3 and 24 hours after rewarming. The maximum endotoxin value for each sample was reported.

#### IEQ estimation

Islets and SC-islets were quantified by DNA content as described in the Supplemental Materials.

#### Endotoxin quantification

Bacterial endotoxin was measured using an Endosafe Nexgen-PTS (Cat. #PTS150K, Charles River). A 100 µL sample of culture supernatant was taken and diluted 10-fold with LAL reagent water (Cat. #NC9276036, Fisher Scientific). The endotoxin level in the diluted sample was measured using the Endosafe analyzer (v.11).

#### Culture sensitivity

A sterile technique was employed during the process. Using a sterile pipette, 100 µL of culture media was dispensed onto the surface of the sterile nutrient agar plate and evenly spread across the plate using a sterile L-shaped spreader. After inoculation, the agar plate was allowed to dry for 5-10 minutes to let the sample absorb into the medium. The plate was then inverted and incubated at 37°C for 5 days. On day 5, the plates were examined for colonies, and colony-forming units were recorded if present.

### Fabrication of CryoMeshes

#### Nylon, stainless steel, and aluminum CryoMesh

CryoMeshes were made using a 3D-printed frame with different mesh sizes and materials. Before fabrication, the metal mesh was cleaned with an acid dip solution (Rio Grande, prepared following the manufacturer’s instructions) for 1 minute and rinsed with DI water for at least 2 minutes. The PLA (Bambu Lab PLC Basic) frame was designed using Autodesk Fusion 360 and 3D printed on a Bambu Lab X1C printer. The nylon, stainless steel, and aluminum CryoMeshes were fused to the frame using a soldering iron at 280°C, controlled by a digital soldering station (RadioShack). The iron’s temperature was within the range of the PLA’s glass transition range. The softened PLA frame penetrated the mesh pores, bonding it to the frame. The CryoMesh was then cleaned with 75% ethanol and DI water sequentially. Stainless steel mesh was sourced from Uxcell, Inc., aluminum from TWP Inc., and nylon from McMaster-Carr, Inc.

#### Gold-coated copper CryoMesh

Copper mesh was obtained from Boegger Industech Ltd. The gold was coated onto the copper mesh via an electroplating process. The copper mesh was cut into the specified sizes (2 × 2 cm, 5 × 4 cm, and 7 × 4.5 cm) and cleaned with 75% ethanol and DI water sequentially before electroplating (described in Supplemental Information).

### Cryoprotective agent preparation

All CPA solutions were prepared in RPMI 1640 medium supplemented with L-glutamine and 25 mM HEPES (Invitrogen Life Technologies, Cat. #22400-089), and filter-sterilized using 0.2 μm nylon membrane filters (Neta-Scientific, Cat. #430049). CPA concentrations are reported as weight/weight percent (wt%).

*CPA Loading Solutions:* Solution 1, 4.4 wt% ethylene glycol (EG, Fisher Scientific), 4.4 wt% dimethyl sulfoxide (DMSO, Fisher Scientific); Solution 2, 11 wt% EG, 11 wt% DMSO; and Solution 3: 22 wt% EG, 22 wt% DMSO.

*CPA Unloading (Rewarming) Solutions:* Solution 4, 11 wt% EG, 11 wt% DMSO, 5 wt% sucrose; Solution 5, 10 wt% sucrose; and Solution 6, 10 wt% sucrose (final dilution step)

### Sterilization techniques for vitrification and rewarming

All equipment, including CryoMesh, CPA solutions, and 3D-printed containers for vitrification and rewarming, was sterilized prior to use. All experiments were conducted in a BSL2 hood to maintain aseptic conditions. CPAs were sterilized using 0.22 μm nylon filters (Cat. #430049, Neta-Scientific). Stainless steel and glassware were autoclaved at 250°F for 30 minutes and dried for 20 minutes before use. Non-autoclavable materials, such as 3D-printed vitrification-rewarming containers and the fabricated *CryoMesh,* were sterilized with an X-Rad 320 (Precision X-Ray, Inc.) at an irradiation distance of 50 cm, operating at 320 kV and 12.5 mA, with a 2 mm aluminum filter to deliver a total dose of 5,000 cGray. All consumables (micropipettes, drapes, RPMI, etc.) were inspected for sterility and expiry date before use.

### Measurement of cooling and rewarming rates

A bare wire type T thermocouple (50 μm diameter, unsheathed; OMEGA) connected to an oscilloscope (DS1M12, USB-Instruments) was used to measure thermal transfer. The thermocouple was fixed at the center of the mesh, where islets were loaded onto different mesh materials and sizes. Cooling and rewarming rates were determined by plunging the CryoMesh directly into liquid nitrogen and subsequently into a rewarming solution. For each measurement, islets were loaded only at the mesh center, and temperature changes from –140°C to –20°C were recorded to calculate rates. The central cooling and rewarming rates were representative of average mesh performance, with spatial variation <5% across the surface (*35*). Coefficients of variation were calculated from repeated measurements for each mesh type.

### Cryopreservation by vitrification using the CryoMesh

During CPA loading, the islets were incubated in Solution 1 (4.4 wt% EG, 4.4 wt% DMSO) for 10 minutes at 21°C, then in Solution 2 (11 wt% EG, 11 wt% DMSO) for 10 minutes at 4°C, and finally in Solution 3 (22 wt% EG, 22 wt% DMSO) for 10 minutes at 4°C. The cluster-CPA suspension was then transferred to the CryoMesh, and excess CPA was wicked off with a sterile Kim wipe. The CryoMesh was then immediately plunged into LN_2_ and stored in a 3D-printed cryo box designed for its specific dimensions.

For rewarming, the CryoMesh was removed from the cryo box while submerged in LN_2_ and quickly immersed in Rewarming Solution 4 (11 wt% EG, 11 wt% DMSO, 5 wt% Sucrose) at 4°C for 10 minutes, followed by *Solution 5* (10 wt% Sucrose) at 4°C for another 10 minutes. Finally, *Solution 6* (with the same composition as *Solution 5*) is used to dilute *Solution 5* for 5 minutes at 21°C, after which only the islets are transferred into their respective culture media and conditions.

### Viability using membrane permeable fluorescent dyes and histology on microscopy

#### Brightfield microscopy

The morphology of fresh and VR islets was captured using a BZ-X800 fluorescence microscope (Keyence).

#### Histology

Fresh and VR islets were fixed in 4% paraformaldehyde and embedded in paraffin within 48 hours. Using a bench-top, room-temperature microtome, 5 μm sections were cut, deparaffinized, and stained with hematoxylin and eosin (H&E), then imaged with a Zeiss microscope. Images were scanned and captured using the ZEN v2.6 (blue edition) software.

### Viability assessment using AO/PI staining

The viability of islets and SC-islets were evaluated using acridine orange (AO) and propidium iodide (PI) membrane-permeability dyes (Millipore Sigma). All reported viability values are relative to control islets or SC-islets for each experiment.

#### Qualitative assessment

Intact islets and SC-islets were stained with 8 ng/mL AO and 20 ng/mL PI for 2 minutes at room temperature. After staining, the samples were coverslipped and imaged using an Olympus Fluoview 3000 inverted confocal microscope equipped with 502/252 nm (AO) and 494/636 nm (PI) filter sets. Images were acquired at a resolution of 4020 × 4020 pixels using 10× or 20× objectives. Note that coverslipping improves depth of imaging but alters the geometry (increased apparent diameter).

#### Quantitative assessment

Three hours after rewarming, islets and SC-islets were enzymatically dissociated into single cells using either TrypLE Express (Thermo Fisher Scientific, Cat. #12605010) or Accutase (Cat. #00-4555-56). Cells were quenched with culture media supplemented with fetal bovine serum and stained with 8 ng/mL AO and 20 ng/mL PI. After 15 seconds of incubation, 10 μL of the cell-dye suspension was loaded onto Countess Cell Counting Chamber Slides (Thermo Fisher Scientific, Cat. #C10228), and viability was measured with the Countess II FL automated cell counter (Invitrogen, Cat. #AMQAF1000).

All qualitative and quantitative findings were validated through 3D reconstruction and image analysis of AO/PI-stained intact islets captured by confocal microscopy.

### Cellular/ mitochondrial respiration and membrane activation

Before assessing mitochondrial respiration, rewarmed islets and SC-islets were recovered at 37°C and 5% CO_2,_ either static in T175 flasks containing culture media for islets or in a dynamic spinner flask at 70 rpm with S3 media for SC-islets.

#### Activated mitochondrial membrane

The islets and SC-islets were stained with 25 μl of 50 ng/ml reconstituted tetramethylrhodamine (TMRE), perchlorate (Biotium, 70005), at room temperature for two minutes, then placed on a microscope slide. A cover slip (Chase Scientific, ZA0294) was added and imaged using an Olympus Fluoview 3000 inverted confocal microscope (excitation/emission at 594/574 nm). Images were captured at a resolution of 4,020 x 4,020 pixels using either a 10X, 20X, or 40X objective. The membrane potential within the islet was quantified by measuring fluorescence intensity with Olympus CellSens Dimension software v4.2.

#### Mitochondrial oxygen consumption

Cellular and mitochondrial respiration were measured using the Agilent Seahorse XF Mito Stress Test and Agilent Seahorse xFe24 Islet Capture FluxPak (Agilent, 103418-100) plates and grids. Islets and SC-islets were handpicked into wells containing 500 μL of culture media, ensuring enough to cover 50% of the inner circle of each well. The islet capture screen was securely fitted onto the grooves within each well’s inner circle. Islets and SC-islets were washed twice with SeaHorse media (SeaHorse XF DMEM) (Agilent, 103575-100), supplemented with 1 mM pyruvate, 2 mM glutamine, and 5.6 mM glucose, then equilibrated for 1 hour at 37°C. Assay reagents were loaded into a pre-hydrated sensor cartridge. The assay plate was inserted into a calibrated Agilent SeaHorse xFe24 analyzer, and the Mito Stress test was performed following the manufacturer’s protocol, using optimized reagent concentrations of 10 μM oligomycin A, 2 μM FCCP, and 10 μM each of rotenone and antimycin A for SC-islets, and 50 μM oligomycin A, 10 μM FCCP, and 10 μM each of rotenone and antimycin A for islets. For normalization between wells, the clusters in each well were lysed with 1 M ammonium hydroxide and 0.2% Triton X-100. DNA content was measured using the PicoGreen assay (Molecular Probes) with standardized calibration controls for quantification on a BioTek Synergy 2 Multi-Mode Microplate Reader (BioTek). Individual cellular respiration parameters, including OCR for basal respiration, ATP production, proton leak, maximal respiration, spare respiratory capacity, and non-mitochondrial respiration, were calculated and normalized to the DNA content of each well.

### Glucose stimulated insulin secretion

Following a three-hour recovery at 37°C with 5% CO2, the samples were either kept static in T175 flasks containing culture media for islets or placed in a dynamic spinner flask at 70 rpm with S3 media for SC-islets. They were then washed once with phosphate buffer saline (PBS) and twice with low-glucose Krebs-Ringer bicarbonate (KRB) buffer containing 3.3 mM glucose. Next, the samples were transferred into 24-well transwell inserts and preincubated in low-glucose KRB for 1 hour at 37°C with gentle rocking to simulate fasting conditions. The islets and SC-islets were washed again with low-glucose KRB and then incubated sequentially in high-glucose KRB with 16.5 mM glucose for one hour, followed by an hour in KRB containing 30 mM KCl, all maintained at 37°C. Supernatants collected after each incubation were stored at -20°C for insulin measurement. After the final step, the islets and SC-islets were dissociated into single cells using TrypLE for 15 minutes at 37°C, and the total cell count was determined with a ViCELL automated cell counter. Insulin concentrations in the supernatants were measured by ELISA (ALPCO, 80-INSHUU-E10) and normalized to the total cell number.

### Transplantation of islets and intraperitoneal glucose tolerance test (IPGTT)

All animal procedures were approved by the Mayo Clinic Institutional Animal Care and Use Committee (IACUC) under protocol A00003973 or by the University of Minnesota IACUC under protocol 1905-37028A. All transplants were performed after a three-hour recovery at 37°C and 5% CO_2,_ either static in T175 flasks with culture media for islets or in a dynamic spinner flask at 70 rpm with S3 media for SC-islets.

#### Immunocompromised non-diabetic xenotransplant model

Male 8- to 10-week-old SCID Beige mice (Charles River Laboratories), weighing 19–24 g, were used as recipients for fresh and VR human islet and SC-islet grafts. Animals were housed in groups in sterile cages with free access to food and water, under controlled ambient temperature (18–25°C), humidity (30–70%), and a 12-hour light/dark cycle. Anesthesia is performed with a 100mg/kg ketamine and 10mg/kg xylazine intraperitoneal injection. Mice were positioned on their right side, and the surgical area was prepped aseptically. An incision was made in the left upper quadrant below the last palpable rib to expose the left kidney by cutting through the abdominal wall. Approximately 3,000 IEQ (∼4.5×10⁶ cells) of islets or SC-islets were transplanted beneath the kidney capsule. The incision was closed in layers, and animals were housed individually and monitored until recovery. At 8 weeks post-transplant, mice were fasted for 6 hours and received a 2 g/kg intraperitoneal (IP) glucose bolus. Blood samples were collected at baseline and 15 minutes post-injection via cheek bleed. Serum was isolated using microvettes (Sarstedt, 20.1292.100), and circulating human insulin was measured by ELISA (ALPCO, 80-INSHUU-E10). The kidney containing the graft was then explanted for imaging.

#### Immunocompromised diabetic xenotransplant model

Male 9–10-week-old NSG (NOD scid gamma) mice, weighing 17-22 g (Jackson Laboratory), were used for transplanting human islets and SC-islets. NSG mice were rendered diabetic via intraperitoneal (IP) injection of 75 mg/kg streptozotocin (Millipore Sigma, S0130) for three consecutive days. Blood glucose (BG) levels were measured after 4-7 days, and diabetes was confirmed by three daily readings exceeding 400 mg/dL. Approximately 4,000 IEQ (6 × 10^6^ cells) of vitrified, rewarmed islets and SC-islets were transplanted under the left kidney capsule following a similar surgical protocol as described above, except anesthesia was induced with 3–4% isoflurane and maintained at 1–2%. BG levels were monitored daily during the first week and then daily on alternating weeks thereafter. Reversal of diabetes and normalization of blood glucose were defined as BG levels below 200 mg/dL on two consecutive days. Graft failure was indicated by two consecutive measurements over 250 mg/dL. On post-operative day (POD) 49, an intraperitoneal glucose tolerance test (IPGTT) was performed after an overnight fast (16 hours) with a 2 g/kg glucose injection dissolved in distilled water. BG levels were recorded every 15 minutes for the first 120 minutes and again at 3 hours. To confirm that the transplanted islets/SC-islets were controlling BG levels and restoring native beta cell function, the left kidney containing the graft was surgically removed, and BG levels were monitored to assess the return of diabetic status and hyperglycemia.

#### Immunocompetent allogenic diabetic model

Female retired breeder Balb/c mice weighing 20-26 g (Charles River Laboratories) were used as islet donors, and 6-8-week-old male C57BL/6 mice (Charles River Laboratories) served as recipients. Islets were isolated from Balb/c mice via collagenase (CIenzyme RI, VitaCyte) digestion and enriched using a Histopaque (Millipore Sigma) density gradient (*23*). The islets were then handpicked and cultured in a bioreactor (ABLE Biott reactor, BWV-S03A) with S3 media for 16 hours before cryopreservation. C57BL/6 mice were rendered diabetic with a single intraperitoneal dose of 200 mg/kg streptozotocin (Millipore Sigma, S0130). Blood glucose levels were monitored after 4-7 days, and diabetes was confirmed by two successive daily measurements exceeding 400 mg/dL. Approximately 500 islets were transplanted under the left kidney capsule of each recipient. No immunosuppressive medication was administered. Blood glucose levels were tracked daily until graft failure, defined as the first of two consecutive measurements exceeding 250 mg/dL.

#### Bioequivalence syngeneic mouse islet transplant

C57BL/6 mice (6- to 8-week-old males, Charles River Laboratories) were made diabetic with a single intraperitoneal dose of 220 mg/kg streptozotocin (Cat. # S0130, Millipore Sigma). Blood glucose levels were measured after 4–7 days, and diabetes was confirmed by two consecutive daily readings exceeding 400 mg/dL. Islet masses of 200 and 150 were compared to a historical dataset of 250 islets per recipient (*23*), transplanted under the left kidney capsule of each recipient mouse. Blood glucose was measured daily. Transplant success was defined as the first of two consecutive days with BG < 200 mg/dL. Graft failure was defined as the first of two consecutive days with measurements exceeding 250 mg/dL.

### Insulin and glucagon immunofluorescence labeling

Explanted left kidneys with the islet and SC-islet grafts were fixed in 4% paraformaldehyde (PFA) overnight at 4 °C. The following day, the grafted kidneys were embedded in paraffin and sectioned. Paraffin-embedded tissue sections were deparaffinized using toluene and rehydrated through a graded ethanol series. Antigen retrieval was performed by boiling the sections in 10 mM sodium citrate buffer (pH 6.0) for 1 hour. After three PBS washes, sections were blocked for 40 minutes in PBS containing 0.1% Triton X-100 and 5% donkey serum. Slides were incubated overnight at 4 °C with primary antibodies against insulin (Cat. #GN-ID4, DSHB) and glucagon (Cat. #G2654-100UL, Sigma-Aldrich). Following incubation, the slides were washed three times (15 minutes each) with PBS and incubated with secondary antibodies Rat Alexa Fluor 488 (Cat. #A21208) and Mouse Alexa Fluor 594 (Cat. #A21203) for 2 hours at room temperature. Sections were then mounted using Fluoromount-G containing DAPI (4′,6-diamidino-2-phenylindole) for nuclear staining. Immunofluorescent images were captured using a Zeiss Axio Observer Microscope with ZEN v2.6 (blue edition) software.

### Apoptosis measurement by quantitative PCR and immunofluorescence imaging

#### RNA isolation and cDNA preparation

Total RNA from control and vitrified islets and SC-islets were isolated after 16 hours in culture using the Arcturus™ PicoPure™ RNA isolation kit (Cat# KIT0204, ThermoFisher Scientific), designed to recover high-quality total RNA. DNase treatment was performed directly within the RNA purification column to remove genomic DNA, following the protocol with the RNase-Free DNase Set (Cat. # 79254, Qiagen). RNA concentration and purity were measured using a NanoDrop 2000 spectrophotometer (ThermoFisher Scientific). Then, 2.0 μg of RNA was used to prepare cDNA with the High-Capacity RNA-to-cDNA™ Kit (ThermoFisher Scientific, Cat# 4387406) according to the manufacturer’s instructions.

#### RT^2^ profiler TM PCR array for human apoptosis and RT-qPCR assay

Real-time PCR was performed using SsoFastTM EvaGreen® Supermix (Bio-Rad Cat #172–5200) on the RT^2^ ProfilerTM PCR Array for Human Apoptosis genes (Qiagen, Cat. #PAHS-012ZD-24 (330231/D-24)) following the manufacturer’s protocol on a Bio-Rad CFX-96 Real-Time PCR Detection System. Fold change in expression between control and vitrified islets/SC-islet was calculated using the ΔΔCt relative expression method. Data analysis and statistical treatment were performed using the vendor-supplied analysis tool (Qiagen).

#### Immunofluorescence labeling for Annexin V

Islets and SC-islets were incubated in a dynamic culture flask at 70 rpm, 37 °C, and 5% CO₂ before the assay. Then, 5× Annexin V binding buffer (Cat. #99902, Biotium) was diluted in deionized water to prepare 1× binding buffer. Clusters were washed twice with 1× binding buffer. The staining solution was made by diluting the Annexin V conjugate in 1× binding buffer to a final concentration of 2.5 μg/mL and incubated with the islets at room temperature for 15–30 minutes, protected from light. The islets and SC-islets were then washed three times with 1× binding buffer and imaged within 30 minutes using the Olympus Fluoview 3000 confocal inverted microscope with an excitation/emission of 490/515 nm.

### RNA sequencing and gene ontology

#### Library preparation and sequencing

Four human islet and four SC-islet preparations were analyzed to evaluate changes in gene expression following VR. Fresh and VR samples were cultured for 16 hours, snap-frozen in liquid nitrogen, and stored at −80 °C until processing. All samples were processed together in a single batch. RNA was extracted using the RNeasy Plus Universal Kit (Qiagen), and TruSeq Stranded mRNA libraries were prepared. The pooled libraries were sequenced on an Aviti Cloudbreak High 2×150 bp run, generating approximately 1 billion pass-filter reads. All expected barcode combinations were successfully detected, with mean quality scores ≥ Q40. Bioinformatic analysis is detailed in Supplementary Materials.

#### Innate immune response by spectral flow cytometry

The *in vitro* IBMIR assay was adapted from prior reports (*57*) and expanded using spectral flow cytometry with 25 markers, as detailed in prior antibody panels (*58, 59*). The study was approved by the University of Minnesota and MHealth Fairview Institutional Review Board (IRB) study #00001261. Type I diabetic (T1D) patients not on immunosuppressive medications were consented before blood was drawn into heparinized vacutainers (Cat. # 366480, BD Biosciences). Transplantation of fresh control and VR islets and SC-islets were simulated by co-culturing T1D whole blood for 1 hour at 37°C and 5% CO_2_ on a rocker set at 22 rpm. At the end of the culture, whole blood was collected, stabilized in Proteomic Stabilizer (PROT1) solution, and stored at -80°C for spectral flow cytometry analysis. Full details of the spectral flow cytometry are in the Supplementary Information.

### Statistical analysis

All statistical analyses were conducted in R (v. 4.5.0; R Foundation for Statistical Computing, Vienna, Austria) unless otherwise noted. The following R packages were used for data processing, statistical analysis, and visualization: tidyverse, edgeR, ggpubr, janitor, ggstatsplot, ggrepel, ggsci, RColorBrewer, ComplexHeatmap, scales, kableExtra, fgsea, circlize, gt, ggforce, rstatix, outliers, superb, ggpval, patchwork, ggh4x, pBrackets, legendry, and magick. Data normality was tested with the Shapiro-Wilk test, and homogeneity of variance was assessed using Levene’s test. Potential outliers were detected using Grubbs’ test. When data met the assumptions of normality and homogeneity, parametric tests were used, such as one-way ANOVA followed by Tukey’s HSD post hoc test. If these assumptions were not met, non-parametric tests such as the Wilcoxon rank-sum test and the Kruskal–Wallis test with suitable post hoc comparisons were employed. Graft survival was compared with the Kaplan-Meier method and the log-rank test. Unless otherwise noted, all tests were two-sided. P values were adjusted for multiple comparisons using the Benjamini-Hochberg method.

## Supplementary Materials

Supplementary Materials and Methods

Supplementary Text

Figs. S1 to S18

Tables S1 to S9

References (*61–88*)

## Acknowledgments

We thank Lifesource Organ Procurement Organization for providing human pancreata, Andy Grams for scientific illustration in Fig. 1, Juan E. Abrahante for processing RNA-seq raw data, and the Minnesota Supercomputing Institute (MSI) for computational resources. We used a AI language models (ChatGPT, OpenAI and Gemini, Google) to assist with drafting brief gene function descriptions, annotating spectral flow cytometry clusters, and refining grammar during manuscript preparation. All AI outputs were reviewed, verified against primary sources, and edited by the authors, who take full responsibility for the final content.

Some figures were created in BioRender (Fig. S13).

## Author contributions

Conceptualization: JSR, ZG, MLE, QPP, EBF, JCB

Methodology: JSR, ZG, MLE, QPP, AH, PR, SR, EBF, JCB

Investigation: JSR, ZG, ZK, KK, DT, SB, PR, AH, MA, AT, BJF, TS, SLB, MS, MLE, SR, QPP, EBF, JCB

Visualization: JSR, ZG, ZK, AH, PR, SR, MLE, QPP, EBF, JCB

Funding acquisition: EBF, JCB, QPP

Project administration: JSR, MLE, EBF, JCB

Supervision: EBF, JCB, QPP

Writing – original draft: JSR, ZG, MLE, EBF, JCB

Writing – review & editing: ZK, KK, DT, SB, PR, AH, MA, AT, BJF, TS, SLB, MS, BNA, BJH, SR, QPP, JSR, ZG, MLE, EBF, JCB

## Data and materials availability

Transcriptomic data are available (NCBI GEO, upload suspended due to the U.S. government shutdown). Raw data for all figures can be found at the Data Repository of the University of Minnesota (DRUM) (*60*).

