## Supplemental Information for "Clinical-grade cryopreservation unlocks transplant-ready human pancreatic and stem cell–derived islets for diabetes therapy"

**The PDF file includes:**

Supplementary Materials and Methods
Supplementary Text
Supplementary Figures S1 to S18
Supplementary Tables S1 to S9
Supplementary References

### Supplementary Materials and Methods

*SC-Islet Media and inductive agents: S1 media* consisted of 1 L MCDB 131 (Life Technologies, 10372019) supplemented with 0.44 g glucose, 2.46 g sodium bicarbonate, 20 µL ITS-X (Life Technologies, 51500-056), 10 mL Glutagro (Corning, 25-015-Cl), 44 mg ascorbic acid, and 10 mL penicillin/streptomycin (P/S) solution (30-001-Cl). *S2 media* consisted of 1 L MCDB 131 supplemented with 0.44 g glucose, 1.23 g sodium bicarbonate, 20 g FAF-BSA, 20 µL ITS-X, 10 mL Glutagro, 44 mg ascorbic acid, and 10mL P/S. *S3 media* included 1L MCDB 131 supplemented with 0.44 g glucose, 1.23 g sodium bicarbonate, 20g FAF-BSA, 5 mL ITS-X, 10 mL Glutagro, 44 mg ascorbic acid, and 10 mL P/S. *BE5 media* consisted of 1 L MCDB 131 supplemented with 3.6 g glucose, 1.754 g sodium bicarbonate, 20 g FAF-BSA, 5mL ITS-X, 10 mL Glutagro, 44 mg ascorbic acid, 10 mL P/S, and 4000 IU heparin (MilliporeSigma, H3149).

KGF (cat # 100-19) was purchased from Peprotech. All other factors were purchased from R&D Systems with the following catalog numbers: Activin A (338-AC), CHIR (4423), SANT-1 (1974), PDBu (4153), Retinoic Acid (RA, 0695), LDN193189 (LDN, 6053), Y27632 (1254), Betacellulin (261-CE), ALK5i (3742), L-3,3',5-Triiodothyronine (T3, 5552), and γ-secretase inhibitor XXI (XXI, 6476).

*IEQ estimation:* Islets and SC-islets were quantified by DNA content using the Quant-iT PicoGreen dsDNA Assay kit (Cat. #P7589, ThermoFisher). Clusters were suspended in culture media with a total volume of 50 mL. After ensuring homogeneity, five aliquots of 100 µL were taken and supplemented with 900 µL of AT buffer (1 M ammonium hydroxide, 0.2% Triton-X 100). The labeled aliquots were sonicated using a Sonic Dismembrator Model 500 (Fisher Scientific) at 20% amplitude for 30 seconds. Aliquots were diluted 100-fold or 400-fold, and a standard curve was prepared in 1X TE buffer with the following DNA concentrations: 500, 250, 125, 62.5, 31.25, 15.625, and 0 ng/mL. 100 µL of each standard and sample was added to separate wells in a 96-well clear-bottom microtiter plate. Standards were assayed in duplicate, while aliquots were assayed in quintuplicates. A working solution was prepared by diluting the assay reagent 200-fold in TE. 100 µL of the working solution was added to each well, and the plate was incubated in the dark for 15 minutes. Fluorescence was measured using a Synergy-2 plate reader (Biotek). A linear regression was applied to the standard measurements, and DNA concentrations of each aliquot were estimated. Using the average from the five aliquots, DNA content was converted to IEQs (10.4 ng DNA/IEQ) and extrapolated to the 50 mL pooled volume of the original preparation to determine the total IEQ.

*Gold electroplating:* The electroplating process is shown in Fig. S18. The electrocleaner solution was prepared by mixing Midas® Electrocleaner Solution Mix (Rio Grande) at 8 oz per gallon in deionized (DI) water and filtering through a 0.22 µm filter (Corning bottle-top vacuum system). The acid dip solution was similarly prepared with Midas Acid Dip Solution Mix (Rio Grande) at 8 oz per gallon in DI water and filtered through a 0.22 µm membrane. The gold plating solution was a ready-to-use Midas Heavy-Deposition 24K Bright Yellow Gold Plating Solution (Rio Grande).

For electroplating, a platinized titanium mesh anode (1.25" wide, Rio Grande) was connected to the negative (–) terminal of a DC power supply (OWON SP3051), and the copper mesh served as the cathode, connected to the positive (+) terminal using an alligator clip.

The process consisted of the following steps:

1. **Electrocleaning:** Copper mesh was submerged in the electrocleaner solution for 1 min at 2.5 V.
2. **Rinsing and acid dip:** The mesh was rinsed in DI water for 1 min, dried with a Kimwipe, immersed in the acid dip for 1 min, and rinsed again in DI water.
3. **Gold plating:** The dried mesh was then immersed in the gold plating solution for 2 min at 2.5 V.
4. **Post-plating rinse:** After plating, the mesh was rinsed sequentially in DI water, acid dip, and DI water, each for 1 min.

The gold-coated mesh was then bonded to a 3D-printed PLA frame, ranging from 2×2 cm to 7×4 cm, with a pore diameter of ~50 µm and wire diameter of ~50 µm, to capture islets and SC-islet clusters (Fig. 2B). To ensure complete surface coverage, a second round of gold plating was performed after bonding. The final assembly was rinsed in 75% ethanol and DI water and prepared for sterilization prior to vitrification. Power was supplied only during the electrocleaning and gold-plating steps.

*RNA-seq Bioinformatics:* RNA-seq data were processed using CHURP (Collection of Hierarchical UMII/RI Pipelines, v1.01) (*61*), developed and maintained by the Research Informatics (RI) group at the Minnesota Supercomputing Institute (MSI), with support from the University of Minnesota Informatics Institute (UMII). CHURP offers a scalable and reproducible framework for RNA-seq analysis, ensuring standardized processing across experiments. The analysis pipeline included:

1. Trimming of raw FASTQ files using Trimmomatic.
2. Quality assessment of raw and trimmed sequences.
3. Alignment of reads to the human reference genome (GRCh38.p13).
4. Generation of QC metrics from BAM alignment files using RNASeQC.
5. Assessment of mapping quality, including concordance and feature counts (HISAT2).
6. Feature quantification using SAMtools.
7. Compilation of a combined gene counts matrix for downstream analysis in R.

*Differential gene expression and downstream analysis:* Differentially expressed genes (DEGs) were identified using the following thresholds: a minimum expression count >50 across all 16 samples, a false discovery rate (FDR) of 7.5% (predefined to increase sensitivity), a nominal p-value <0.05, and an absolute $\log_2$ fold change >1. The analysis workflow included:

1. Unsupervised clustering of the 500 most variable genes across all samples.
2. Paired differential expression analysis (control vs. VR) for each islet donor or SC-islet preparation using the Genewise Negative Binomial Generalized Linear Model with Quasi-likelihood Tests (glmQLFit, edgeR).
3. Identification of DEGs based on FDR, p-value, and $\log_2$ fold change criteria.
4. Pathway enrichment analysis using Over-Representation Analysis (ORA) with the Hallmark Molecular Signature gene set (*62*), implemented via the clusterProfiler (*63*) package in R. Genes with absolute $\log_2$ fold change >1 and p-value <0.05 were included.

Enriched pathways were filtered using an adjusted p-value <0.01 and q-value <0.2. Additionally, Gene Set Enrichment Analysis (GSEA) was performed using the ShinyGO online platform (*64*) to assess enrichment within Gene Ontology (GO) pathways. The analysis used annotations from the curated PANTHER database (*65*), applying an FDR threshold of 0.01.

*Spectral Flow Cytometry:*

*Staining of innate immune cells:* All staining was performed in standard FACS tubes, using $1.0 \times 10^6$ PBMCs for single-color tests and $2.0 \times 10^6$ PBMCs for multicolor tests. Reference control cells were centrifuged at 1000xg for 5 minutes at room temperature (RT) to remove media. Blood samples stabilized in PROT1 were thawed at 4°C. Red blood cells were lysed with ACK lysis buffer (Cat. #A1049201, Thermo Fisher). The cell suspensions were washed twice with FACS buffer. Cell pellets were resuspended and washed with 200 µl of PBS.

To evaluate viability, 100 µL of ViaDye Red Viability dye was added to both the control and test samples and incubated for 20 minutes at room temperature. Meanwhile, 100 µL of staining buffer was added to the remaining single-color control and unstained tubes. After incubation, all tubes were centrifuged, the supernatants discarded, and the cells washed with 200 µL of staining buffer. A second centrifugation was performed, and the supernatants aspirated. Each single-color control was stained with its respective antibody, while the test samples were incubated with the complete antibody cocktail (prepared in 200 µL with 10 µL of Brilliant Staining Buffer Plus (Cat. #AB_2869761, BD Biosciences)). All antibodies were pre-centrifuged to remove aggregates that could affect staining quality. Unstained tubes received only staining buffer. After a 30-minute incubation at 4°C, the samples were washed and centrifuged to remove unbound antibodies. A final rinse with 200 µL of staining buffer was performed before fixation. The cells were fixed in 2% paraformaldehyde for 20 minutes at room temperature. A subsequent wash step ensured the removal of residual PFA.

*Data acquisition:* Stained cells were resuspended in 300 µl of staining buffer, and data were acquired using a five-laser Cytek Aurora flow cytometer system (355, 405, 488, 561, and 640 nm) with custom-designed and validated antibody panels. This process involved staining for 25 key surface markers to assess neutrophils, monocytes, dendritic cells, natural killer cells, NKT cells, and myeloid-derived suppressor cells.

*Spectral unmixing*: The optimal fluorochrome combinations for the panel were initially evaluated using Cytek Bioscience's similarity index, which ranges from 0 (completely different) to 1 (identical). Fluorochromes with a similarity index of 0.98 or below were deemed suitable for combined use. The compatibility of these fluorochrome pairs was further assessed with Cytek's complexity index, which considers spectral overlap, spillover, and autofluorescence effects. Lower complexity scores indicate better performance with less signal spillover. Using these metrics, 25 fluorochromes were selected to identify the frequencies of monocyte, neutrophil, dendritic cell, natural killer cell, and myeloid-derived suppressor cell (MDSC) subsets. The full antibody panel are listed below.

*Data analysis:* For each cell type group, transformed channel values were exported from FlowJo (Becton, Dickinson & Company) as CSV files, then imported and processed using the scanpy (v1.11.2) package (*66*) in Python (v3.11.12). Briefly, we first performed initial dimensionality reduction with principal components analysis (PCA), followed by nearest-neighbor distance and graph inference, then applied UMAP (Uniform Manifold Approximation and Projection) for further dimensionality reduction. Clusters were inferred from the neighborhood graph across a range of resolutions (0.1-0.9) using the Leiden algorithm (*67*). The complete data set for each group was then exported in anndata format (v0.11.4) and imported into R (v4.5.0) for further analysis.

We performed bootstrap resampling (5,000 iterations) using the boot (v1.3-31) package to estimate group-wise statistics, correct for potential bias, and generate nonparametric 95% confidence intervals for comparisons (*68*). This nonparametric approach enabled robust inference without assuming normality and provided a measure of uncertainty around group differences based on the resampled distribution of the test statistic. For visualization, UMAP plots were downsampled (tidytof v. 0.99.8 (*69*)) to ensure equal cell numbers within each parent gate (e.g., neutrophils), providing balanced representation across all subgroups. Expression matrices were scaled by cluster and marker to demonstrate relative expression across clusters and displayed as heatmaps (tidytof). Cluster phenotypes were assigned using an AI-assisted (ChatGPT and Gemini) algorithm with manual supervision using scaled and unscaled fluorescence intensities.

*Antibodies used in spectral flow cytometry:*

| Antibody | Fluorophore | Vendor | Catalog# | Clone |
|---|---|---|---|---|
| CD45 | B548 | Cytek | R7-20291 | HI30 |
| CD56 | BV785 | BD Biosciences | 564058 | NCAM 16.2 |
| CD3 | PE-Fire640 | BioLegend | 344860 | SK7 |
| CD19 | BYG710 | Cytek | R7-20009 | HIB19 |
| CD14 | PECy7 | BD Biosciences | 557742 | 9095611 |
| CD7 | BV480 | BD Biosciences | 566119 | M – T701 |
| CD16 | BV570 | BioLegend | 302036 | 3G8 |
| HLA-DR | R840 | Cytek | SKU R7-20294 | L243 |
| CD33 | PE | BD Biosciences | 555450 | 1165676 |
| CD11b | B515 | BD Biosciences | 564517 | ICRF 44 |
| CD20 | B675 | Cytek | R7–20195 | 2H7 |
| CD5 | BV510 | Bio legend | 364018 | 017F12 |
| CD141 | BV750 | BD Biosciences | 747244 | 1A4 |
| CD1c | BV650 | BD Biosciences | 569617 | F 10/21A3 |
| CD123 | PECy7 | BD Biosciences | 551065 | 8250742 |
| Live/Dead | ViaDye™ Red | Cytek | SKU R7-60008 | - |
| CD54 | BUV496 | Bio legend | 353126 | B297158 |
| CD184 | BUV395 | BD Biosciences | 740265 | B11/CXCR4 |
| CD49d | BUV737 | BD Biosciences | 748702 | L25 |
| CD63 | PerCP-Cy5.5 | Bio legend | 353020 | 85C6 |
| CD15 | APC | Bio legend | 323008 | B421788 |
| OLFM-4 | AF488 | Novus bio-techne | 06415AF488 | D185256 |
| CD66b | BV421 | Bio legend | 392916 | 412789 |
| CD62L | PE-Dazzle594 | Bio legend | 304842 | B418094 |
| CD177 | BUV615 | BD Biosciences | 751453 | 507-3600 |

### Supplementary Text

#### *RNA-seq*

Overall, the transcriptomic changes associated with VR were minimal. For islets, there were no statistically significant changes (Fig. 5A, Table S3). For SC-islets, 20 genes were upregulated (Fig. 5B, Table S4), but the magnitude of the change was small ($\log_2 FC \leq 5.3$). DGE transcripts were linked to cell stress, apoptosis, and NF-κB pathways (Table S5). Among the upregulated genes, **PPP1R15A (GADD34)** regulates stress responses via the unfolded protein response and promotes apoptosis during prolonged stress (*70*). **GADD45A** is involved in cell cycle arrest and apoptosis following genotoxic stress or DNA damage (*71*). **HSPA7**, a pseudogene member of the heat shock protein family transcribed in response to cell stress but not producing a functional protein (*72*). **TNFRSF10D**, a decoy receptor for TRAIL, inhibits apoptosis by blocking death signaling cascades (*73*). **SQSTM1 (p62)** acts as a multifunctional adaptor in oxidative stress, autophagy, antioxidant defense, and NF-κB signaling (*74*). **NFKBIZ** modulates NF-κB activity and the inflammatory stress response (*75*). **ATF3** is a transcription factor activated by cellular stress and DNA damage (*76*). **TNFAIP3 (A20)** is a key regulator of inflammation and apoptosis, serving as a negative feedback inhibitor of NF-κB, and is also involved in autophagy and stress resolution (*77*).

Compared to a single-cell RNA-seq study of fresh and cryopreserved islets (*53*) that reported many DGE, we found no overlap except for two PP1 regulatory subunits: **PPP1R1A**, a PP1 inhibitor responsive to cAMP signaling, was downregulated in their study, while **PPP1R15A/ GADD34**, which modulates PP1 during ER stress, was upregulated in ours.

### Supplementary Figures:

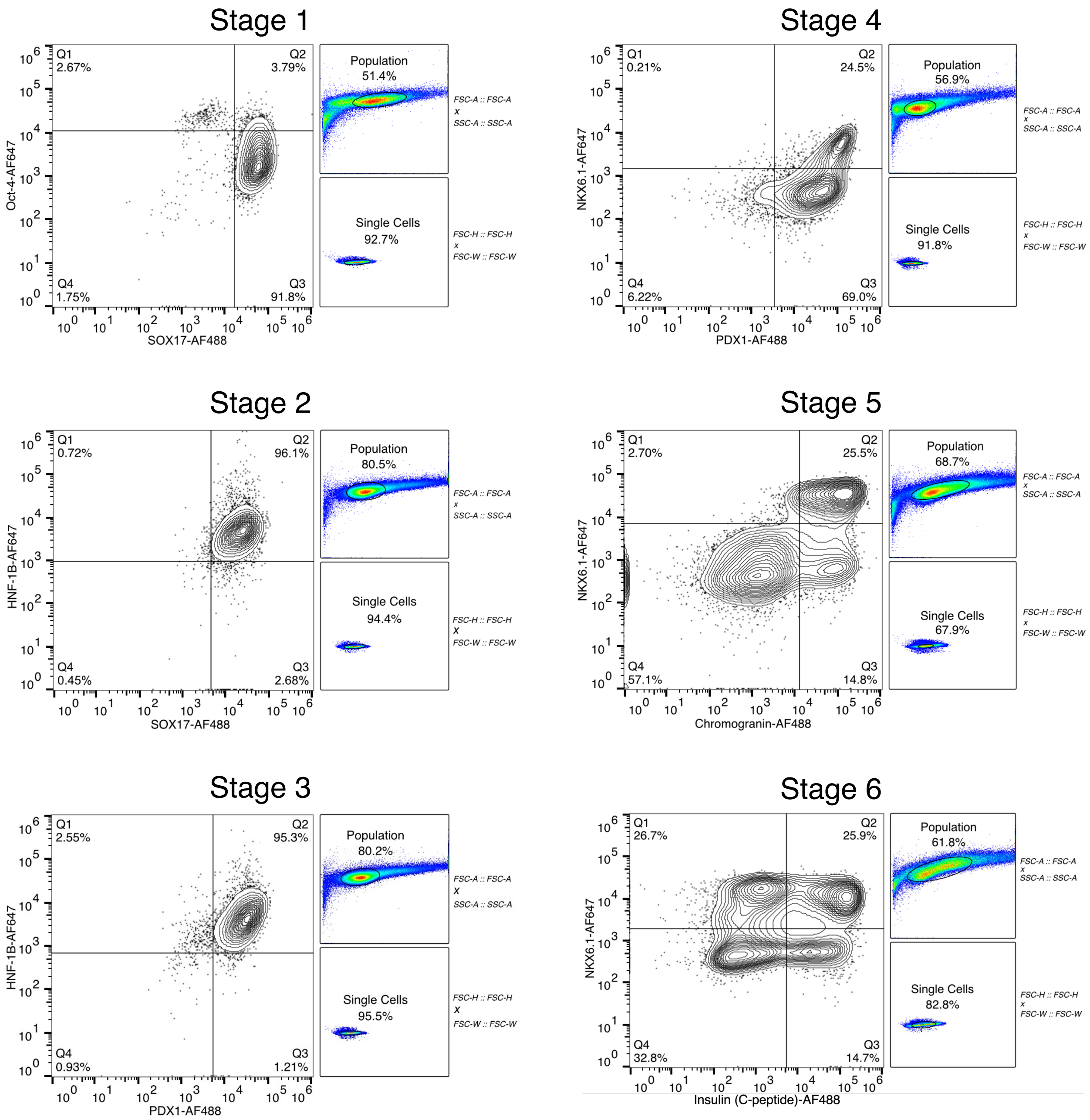


**Fig. S1. Characterization of each stage of SC-islet differentiation**. HUES-8 human embryonic stem cells were differentiated through stages by sequentially changing the growth factor-containing media. SC-islets were dissociated and analyzed by flow cytometry using indicated markers. Anti-insulin antibody is against C-peptide portion.

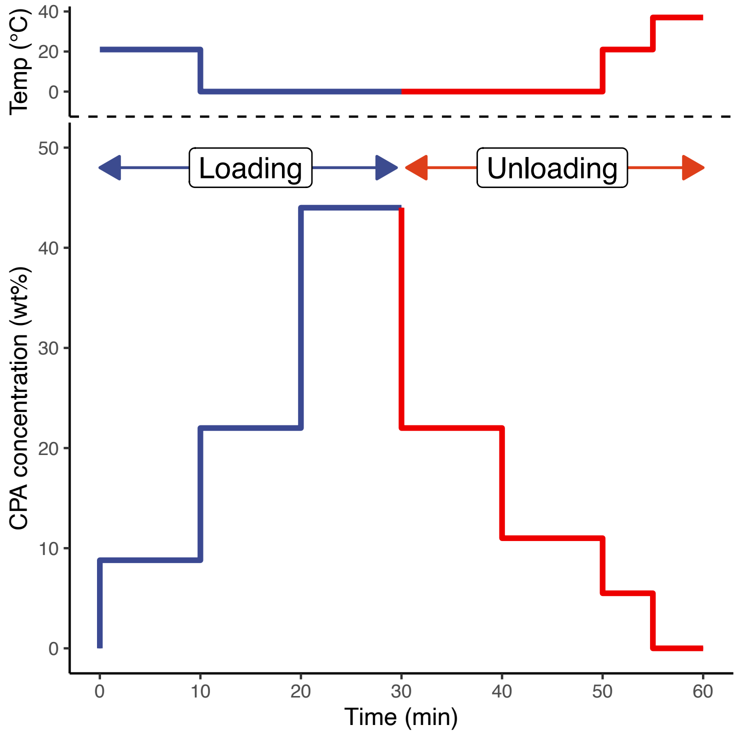


**Fig. S2. Stepwise cryoprotective agent (CPA) loading and unloading protocol**. The CPA concentration and temperature profiles are shown (*23*).

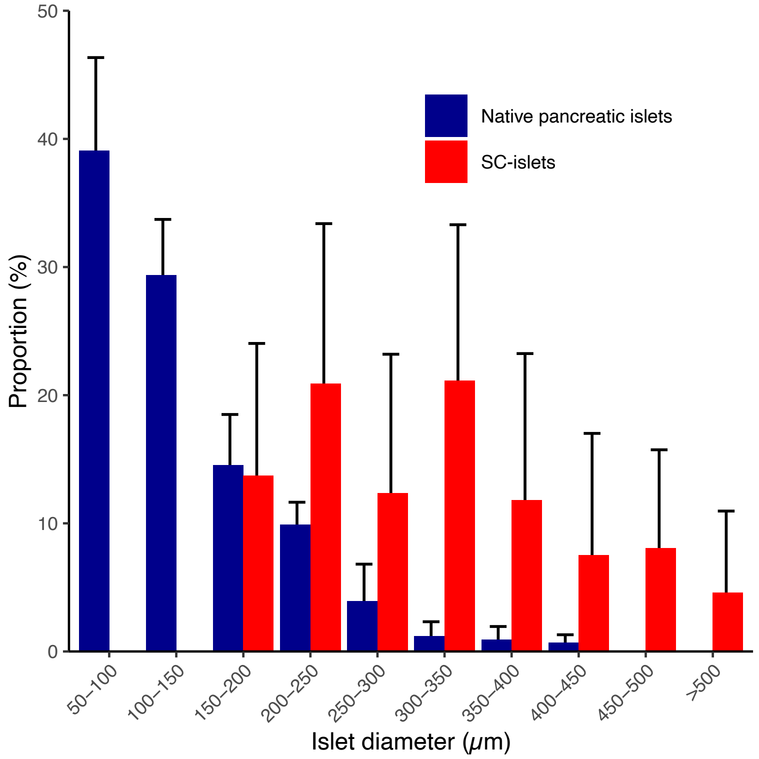


**Fig. S3. Human pancreatic islet and SC-islet size distribution (diameter).** Mean ± SD.

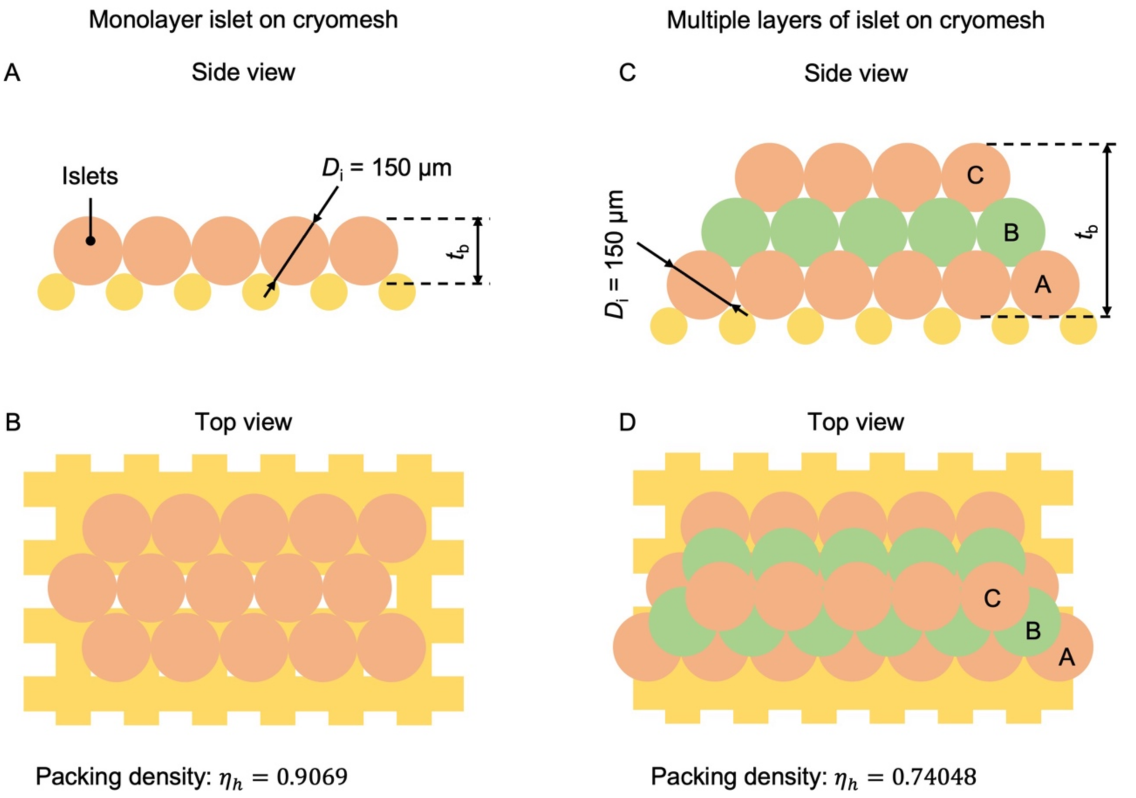


**Fig. S4. Theoretical islet density on CryoMesh**. Schematic of the side view (**A**) and top view (**B**) of monolayer islets on the CryoMesh. Schematic of the side view (**C**) and top view (**D**) of multiple layers of islets on the CryoMesh. The schematic and calculation are based on Hales (*78*).

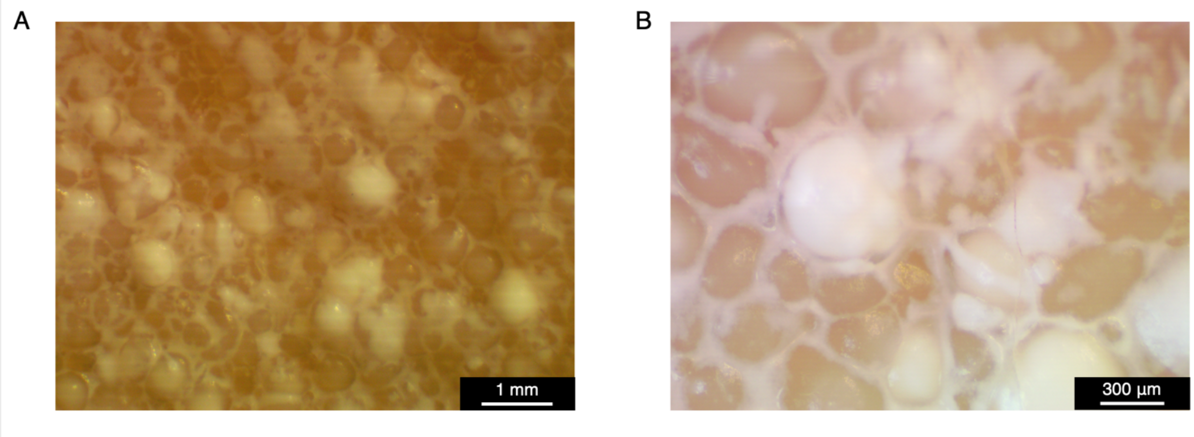


**Fig. S5. Ice formation in islets with a diameter > 500 µm.** Magnified images of islets in CPA after CryoMesh cooling. (**A**) Macro view with a scale bar of 1 mm. (**B**) Micro view with a scale bar of 300 µm. Opaque appearance indicated ice formation. Vitrified islets remain translucent (Fig. 2C).

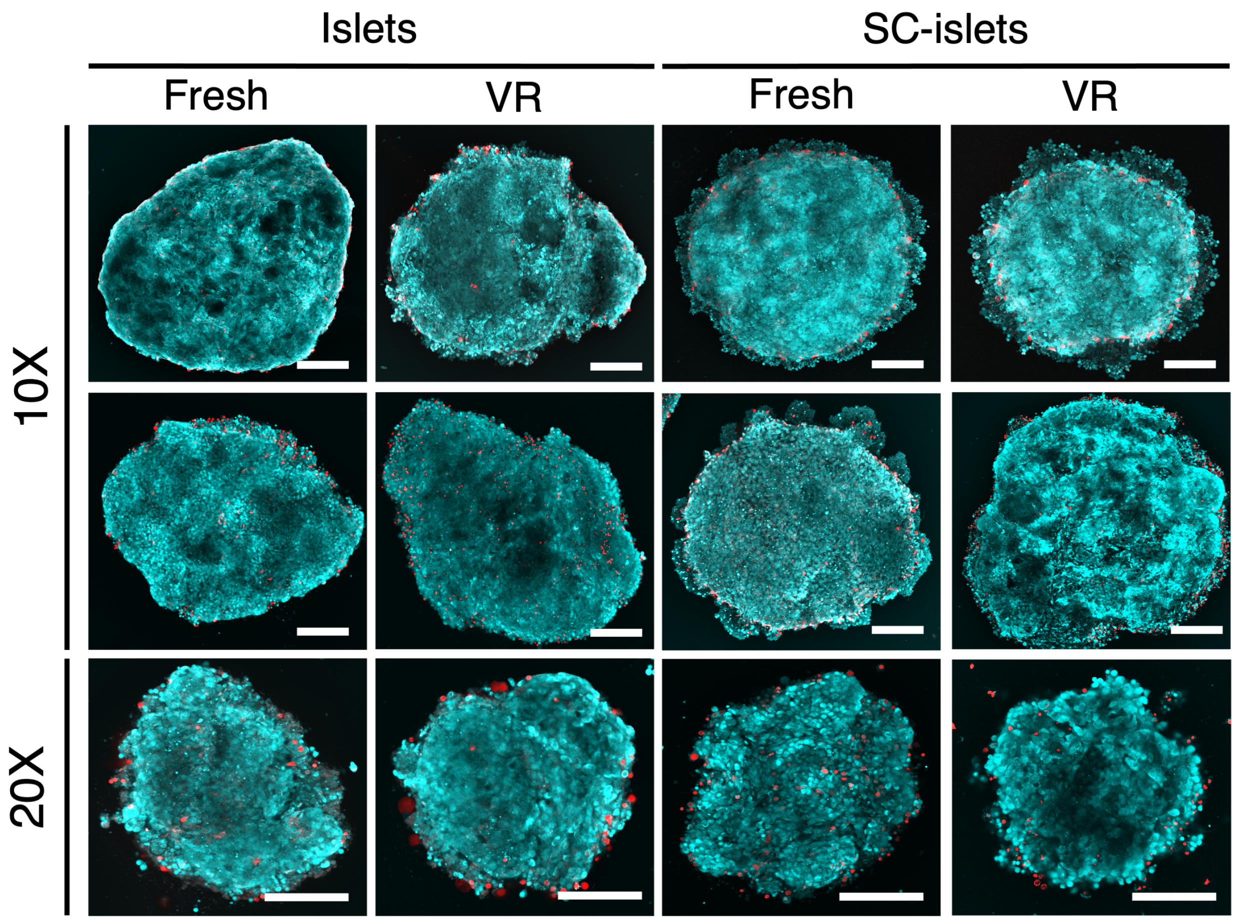


**Fig. S6. Additional viability imaging of islets and SC-islets after VR.** Additional confocal images from viability imaging with acridine orange (teal, live cells) and propidium iodide (red, dead cells). Sample images at different magnifications are shown. Scale bars set at 75 μm.

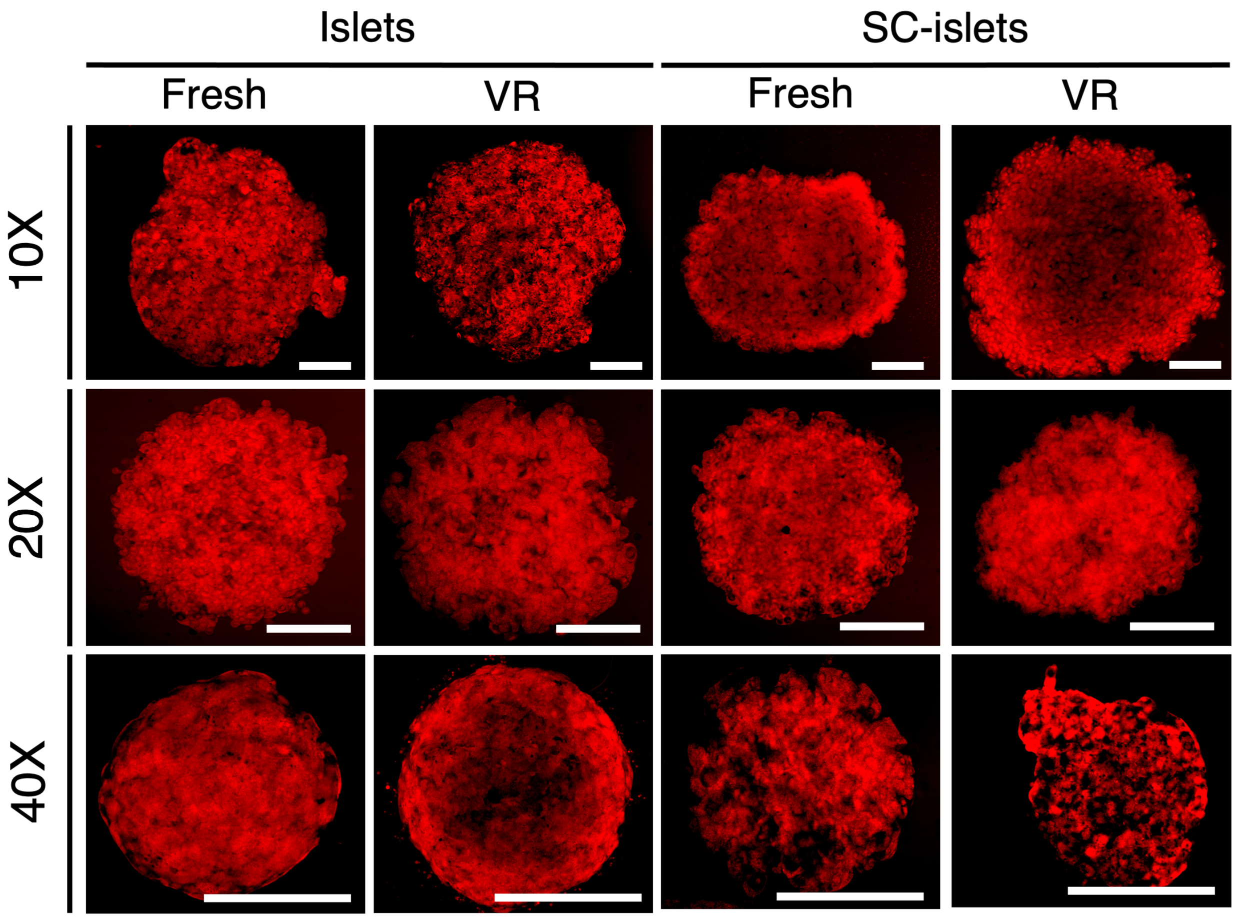


**Fig. S7. Additional imaging mitochondrial membrane potential.** Additional confocal images of tetramethylrhodamine ethyl ester (TMRE)-stained (red) islets or SC-islets showing preserved mitochondrial membrane potential. Sample images at different magnifications are shown. Scale bars set at 75 μm.

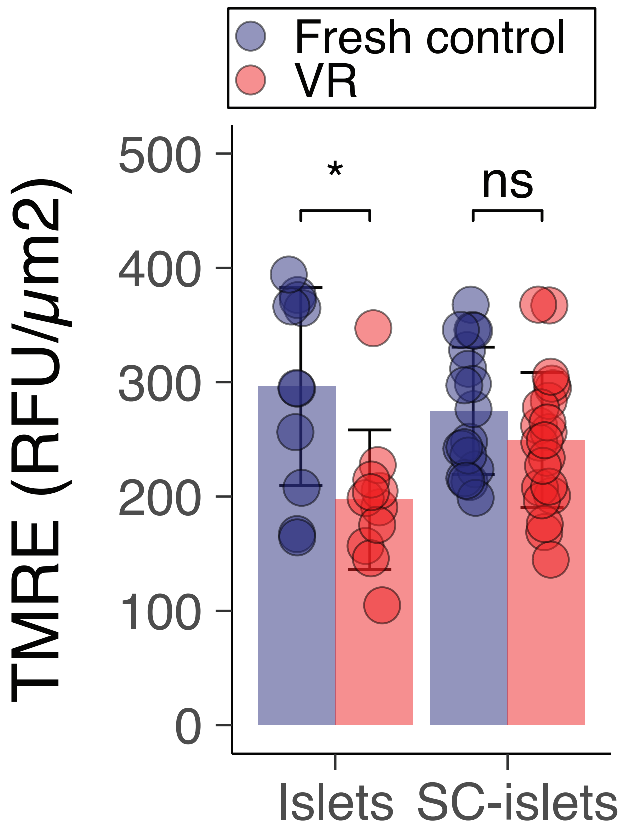


**Fig. S8. Quantification of mitochondrial membrane potential.** Image analysis of confocal images of tetramethylrhodamine ethyl ester (TMRE)-stained islets or SC-islets was performed to quantify fluorescence intensity at 3 hours post-rewarming. We previously showed TMRE staining improves further at 24 hours (*23*). Mean ± SD, T-test, *, $p < 0.05$. ns, non-signicant; RFU, relative fluorescence units; SC, stem cell; VR, vitrified and rewarmed.

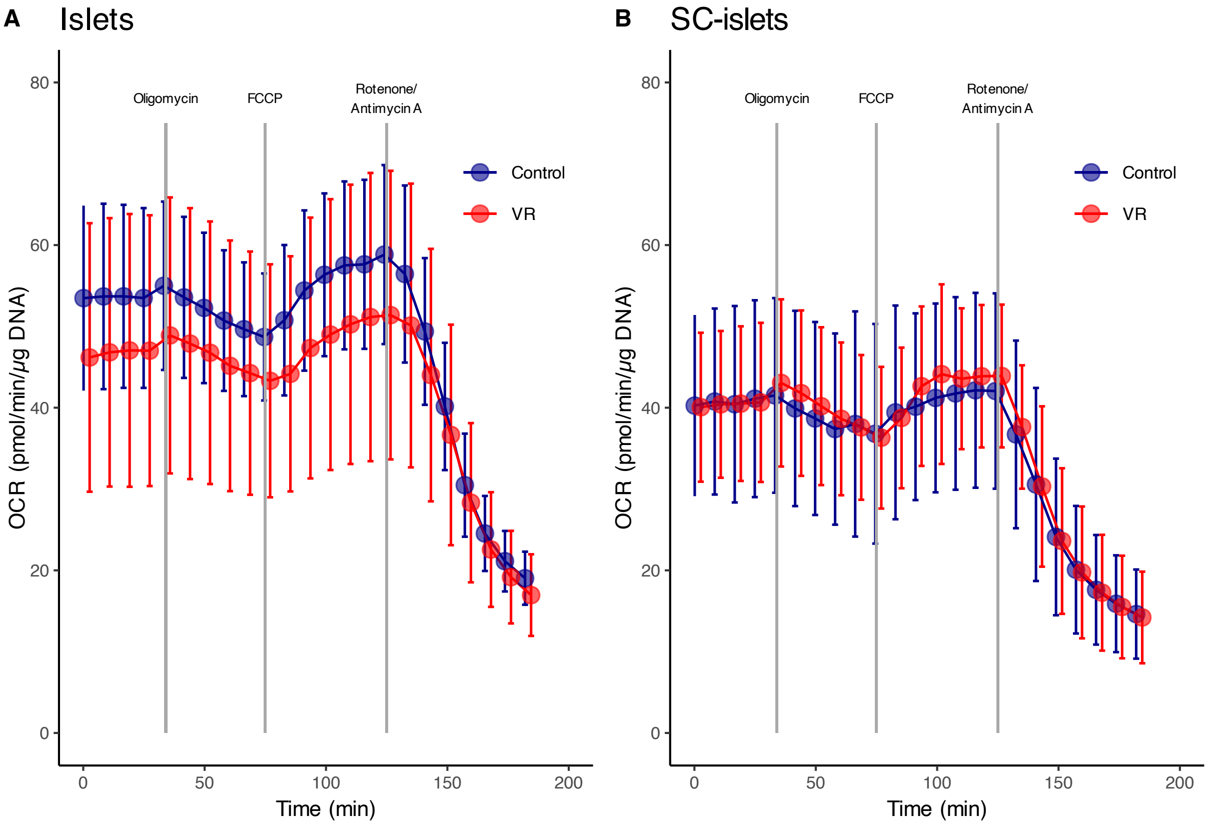


**Fig. S9.** Mitochondrial stress test of oxygen consumption in fresh and vitrified/rewarmed (VR) samples. Oxygen consumption rate (OCR) was measured using a Seahorse mitochondrial stress test following sequential treatment with oligomycin, FCCP, and rotenone/antimycin A. (**A**) Human islets. All p = ns, Kruskal-Wallis test and Wilcoxon post-hoc test, n = 71 per timepoint; and (**B**) SC-islets. All p = ns, Kruskal-Wallis test and Wilcoxon post-hoc test, n = 24-26 per group and per timepoint. Fresh samples are indicated in red, and VR samples in blue. FCCP, carbonyl cyanide-4-(trifluoromethoxy)phenylhydrazone; ns, not significant; VR, vitrified and rewarmed.

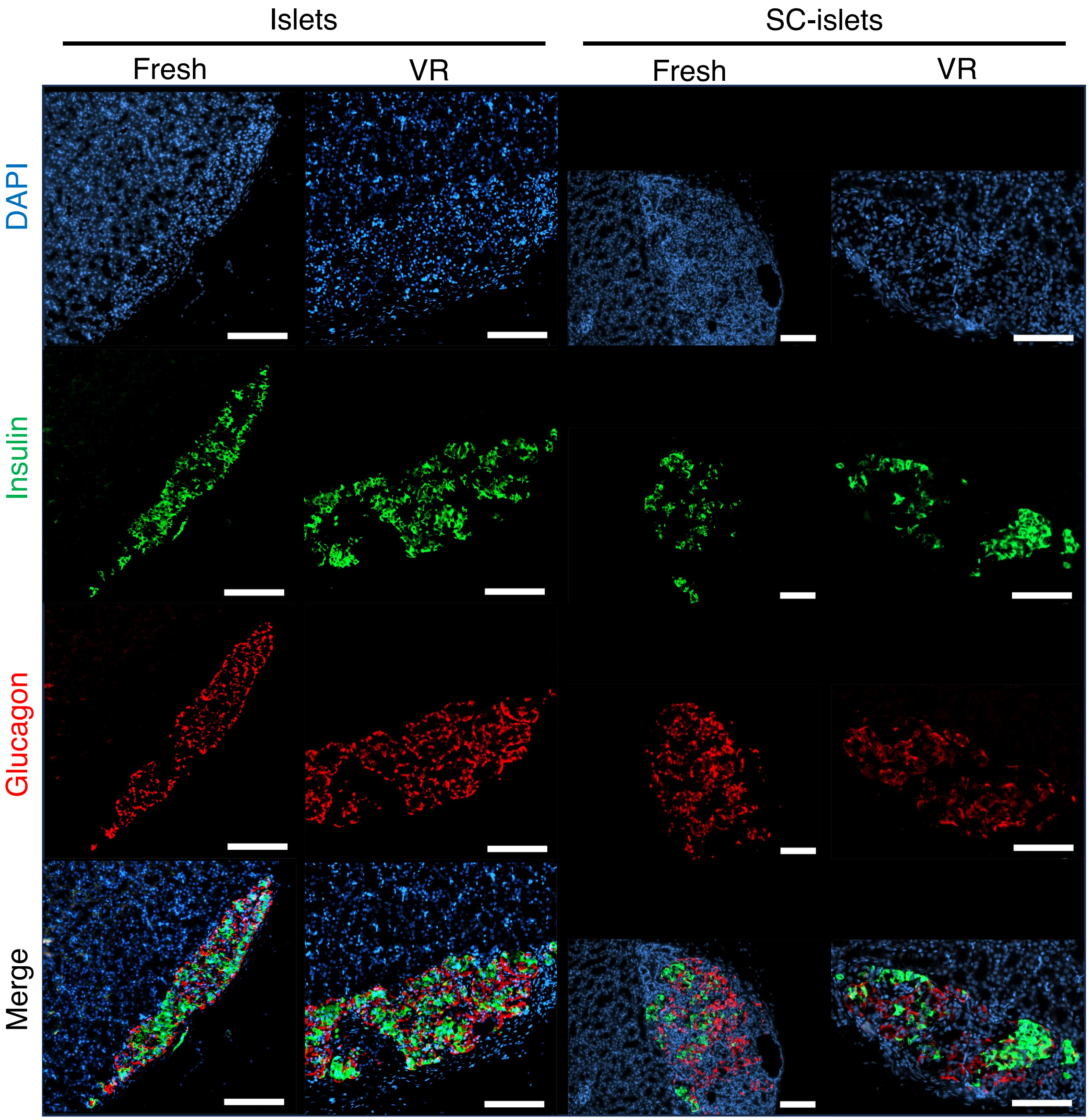


**Fig. S10.** Confocal imaging of transplanted human islets and SC-islets. Native human pancreatic islets or SC-islets were transplanted under the kidney capsule of immunodeficient SCID-Beige mice. At 12 weeks posttransplant, grafts were explanted and sectioned for immunofluorescence staining of insulin (green), glucagon (red), and nuclei (DAPI, blue). Individual channels and merged images are shown. Scale bar set at 100 μm.

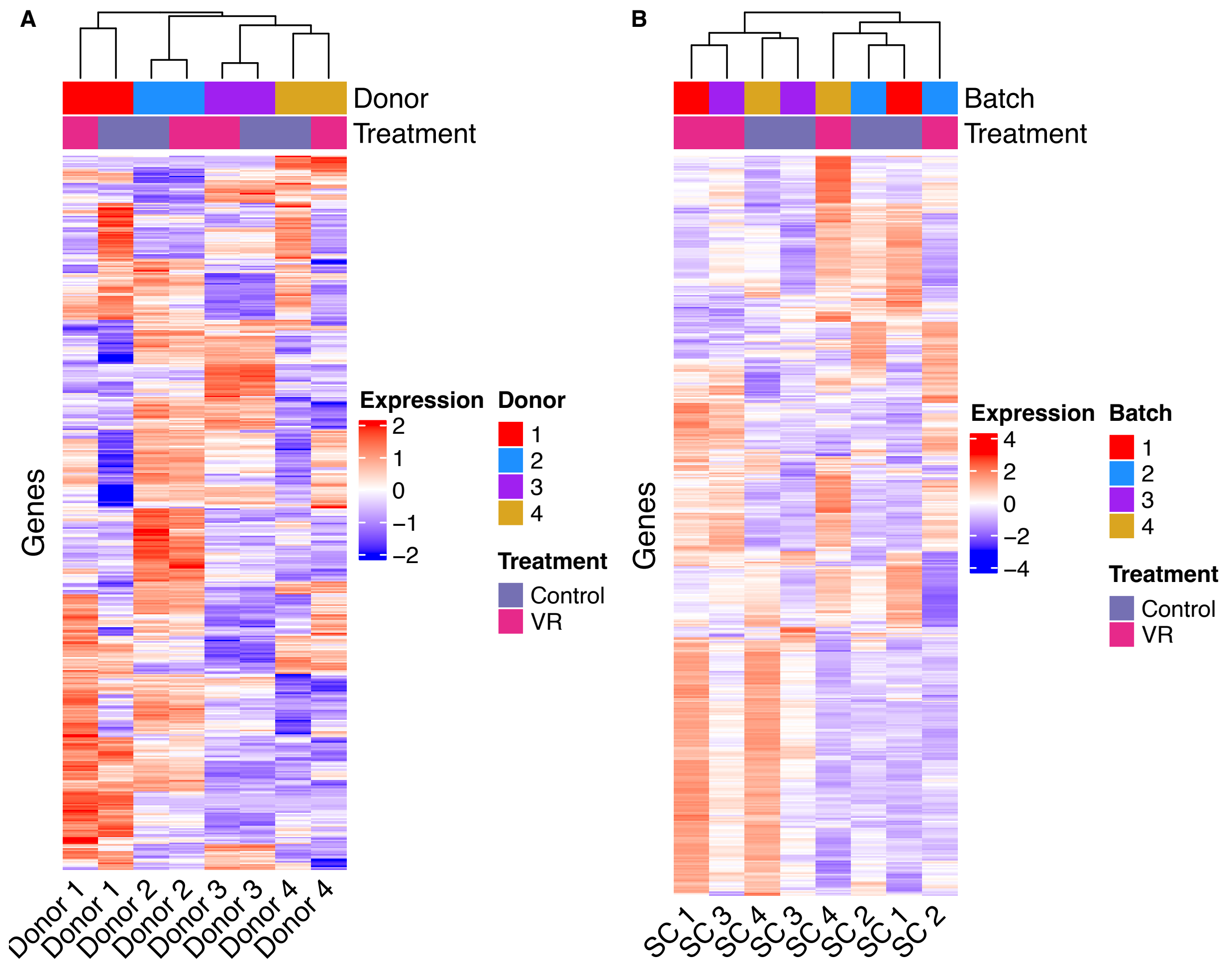


**Fig. S11.** RNA-seq analysis of fresh versus vitrified/rewarmed (VR) samples. (A) Native human islets and (B) SC-islets were analyzed by RNA sequencing. Relative gene expression profiles were compared between fresh and VR samples from four islet donors and four independent SC-islet batches. Heatmaps display unsupervised clustering of the 500 most differentially expressed genes. Clustering segregates samples both by donor/batch and by treatment condition (VR v. fresh).

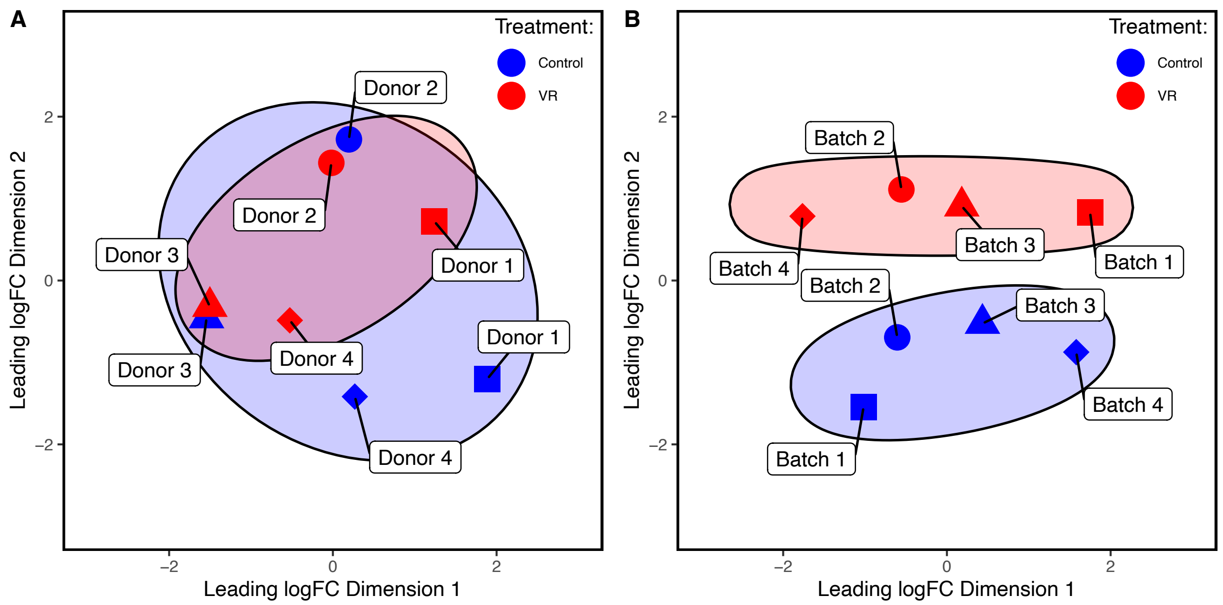


**Fig. S12.** Multidimensional scaling (MDS) plots of RNA-seq comparing fresh and VR islets SC-islets. (A) Pancreatic islets and (B) SC-islets. Each point represents a sample from one of four islet donors or SC-islet batches. Control (fresh) islets and clusters are shown in blue, while vitrified and rewarmed (VR) islets and clusters are in red. Ellipses indicate group clustering, showing the extent of overlap or separation between control and VR samples.

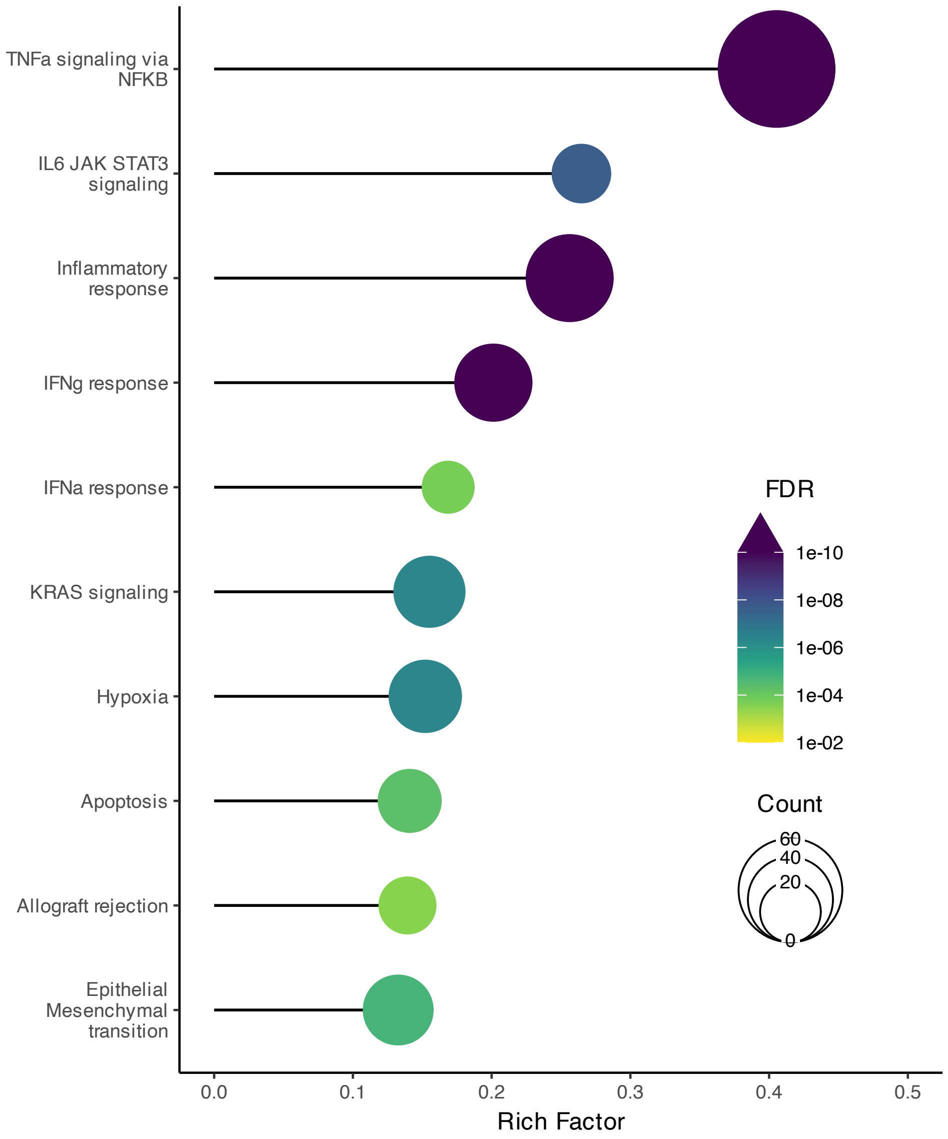


**Fig. S13. Gene set enrichment analysis (GSEA) of RNA-seq data comparing fresh and vitrified/rewarmed (VR) SC-islets using the hallmark gene ontology database.** Results are shown as a lollipop plot, where bar length represents the Rich Factor (ratio of differentially expressed to total genes in a pathway). Circle size indicates the number of genes in each pathway, and the color gradient of the circles shows the adjusted false discovery rate (FDR), with darker colors indicating stronger statistical significance.

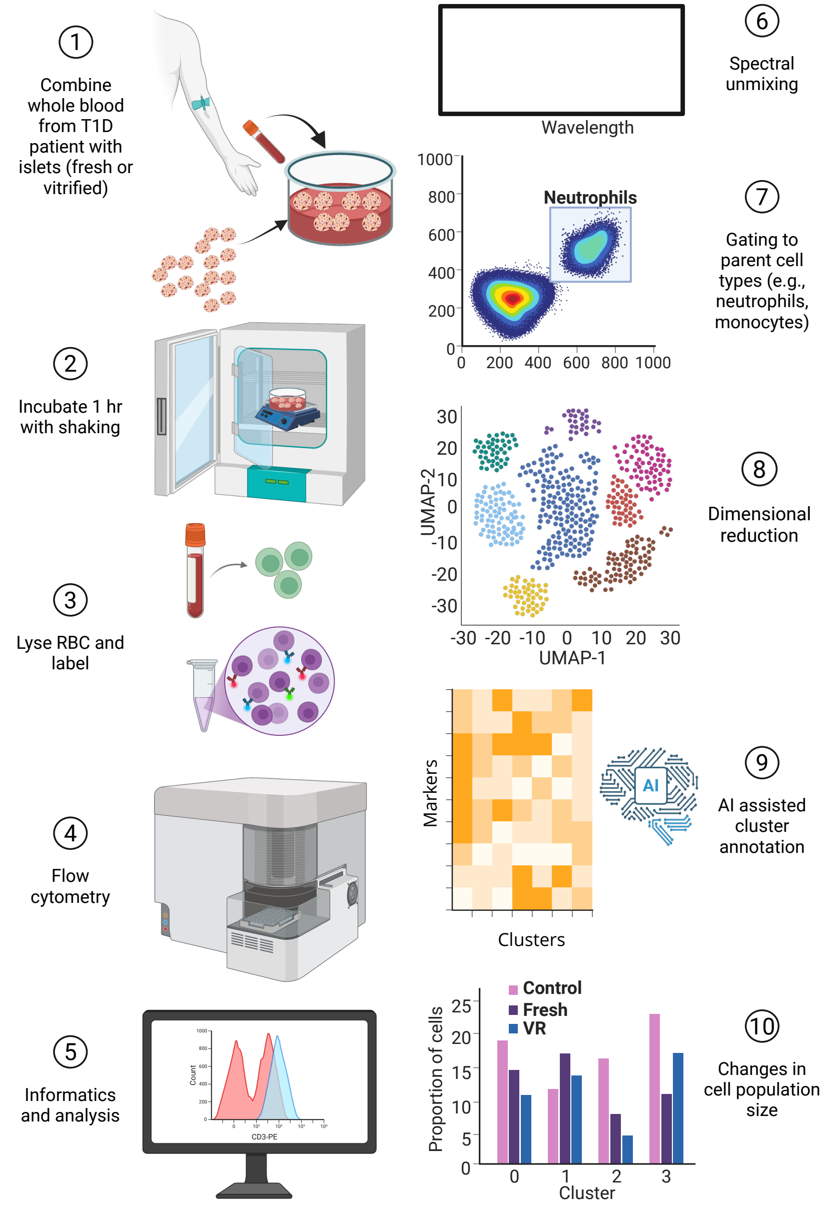


**Fig. S14.** *In vitro* assay of innate immune activation simulating immediate blood-mediated inflammatory response (IBMIR). The workflow proceeds as follows: (1) fresh or vitrified/rewarmed (VR) islets or SC-islets were mixed with whole blood from type 1 diabetic patients; (2) mixtures were incubated at 37 °C for 1 hour; (3) red blood cells were lysed, and leukocytes labeled with a 25-parameter fluorescent antibody panel; (4) spectral flow cytometry was performed; (5) computational informatics applied; (6) spectral unmixing carried out; (7) standard gating used to identify parent cell populations (neutrophils, monocytes, dendritic cells, MSDCs, NK/NKT/lymphocytes); (8) dimensional reduction applied with UMAP; (9) cluster phenotypes assigned using AI-assisted comparison of relative marker fluorescence intensities; and (10) relative subpopulation sizes compared across conditions (blood alone, blood + fresh islets/SC-islets, or blood + VR islets/SC-islets). Created in BioRender. AI, artificial intelligence.

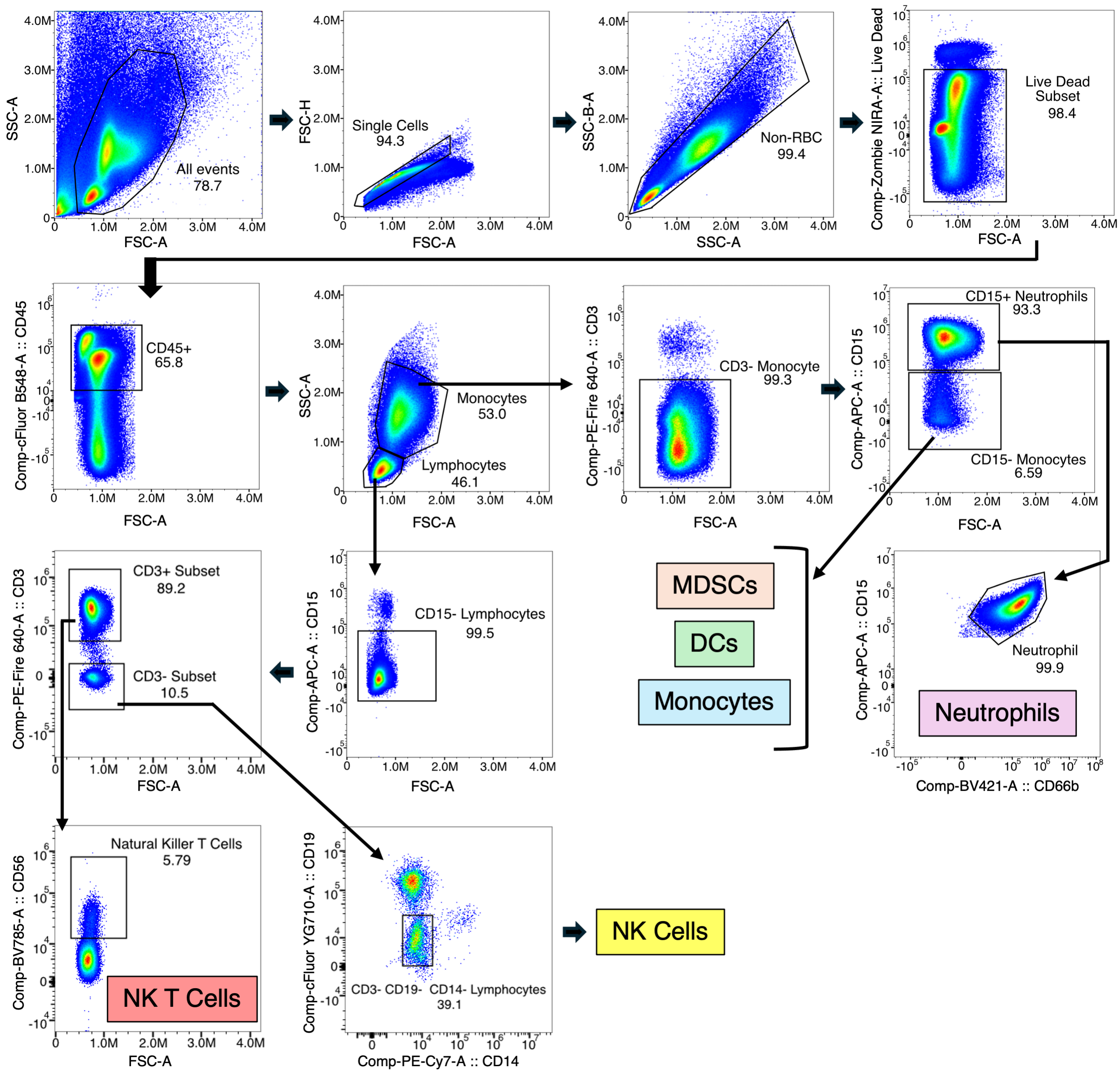


**Fig. S15. Innate flow cytometry of parent cell types.** The diagram illustrates gating strategies to identify the neutrophil, monocyte, dendritic cells, myeloid-derived suppressor cells, and NK/NKT cell parent populations for subsequent analysis.

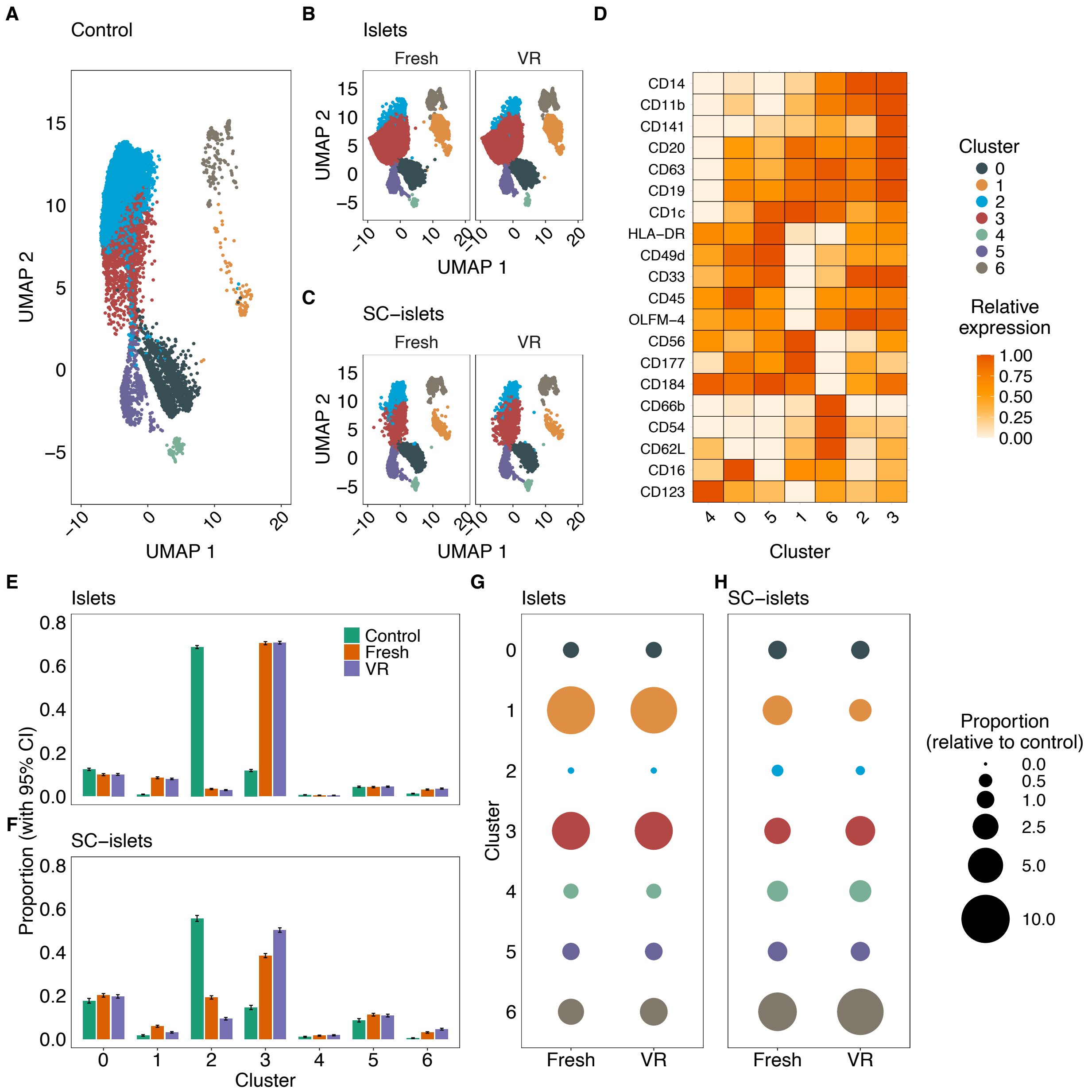


**Fig. S16. Instant blood-mediated immune responses (IBMIR) in monocytes, dendritic cells, and myeloid-derived suppressor cells (MDSC).** Dimensional reduction analysis depicting cell subpopulation clusters for (**A**) control blood samples, (**B**) fresh and vitrified/rewarmed (VR) islets, and (**C**) fresh and VR SC-islets. (**D**) Transformed and scaled relative expression of key cell surface markers identified in **A-C**. Population sizes for control blood samples, fresh islets/ SC-islets, and VR islets/SC-islets are shown for each cluster in (**E**) islets and (**F**) SC-islets. Data are mean ± 95% confidence interval. Bubble plots illustrate the relative population size compared to control for (**G**) fresh and VR islets and (**H**) fresh and VR SC-islets.

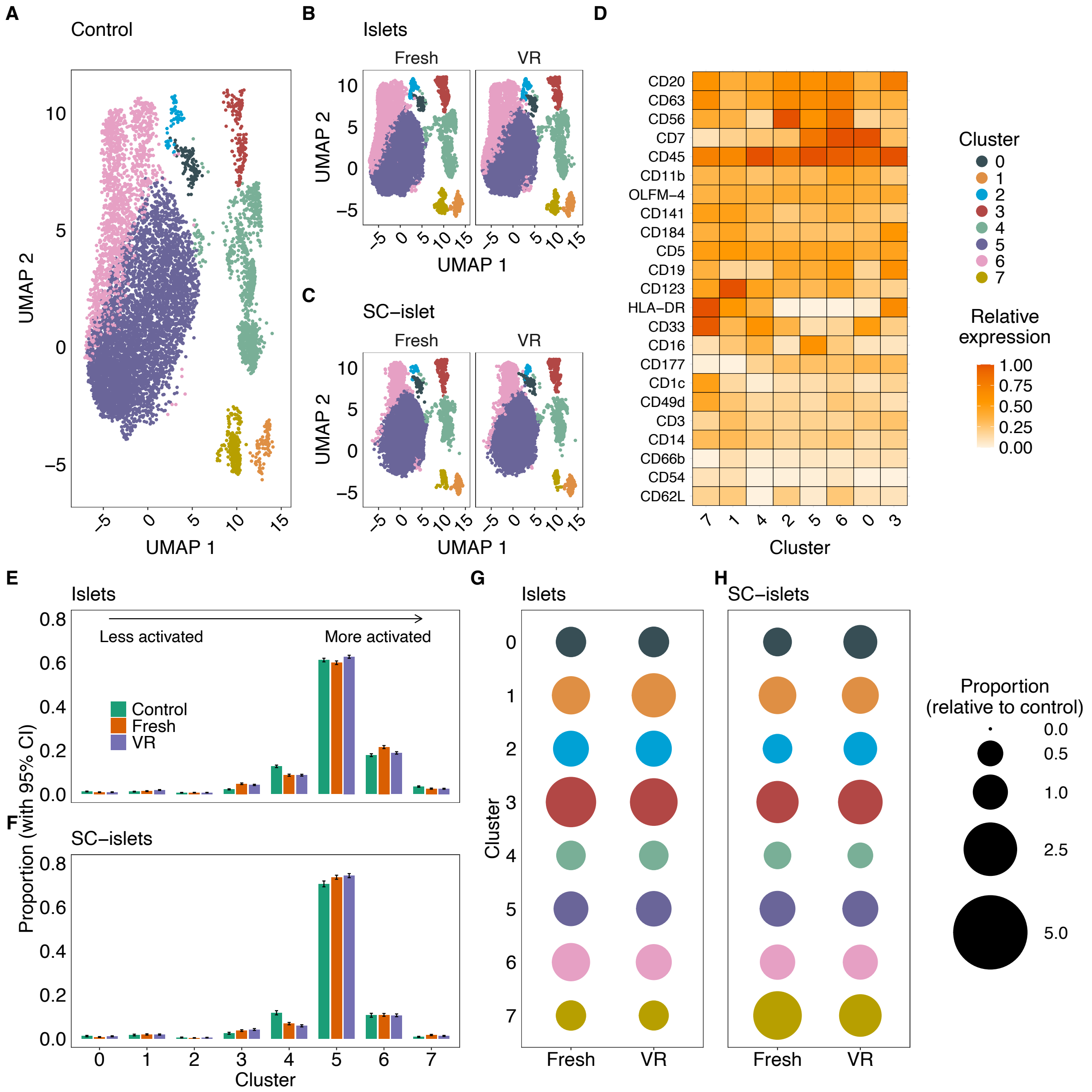


**Fig. S17. Instant blood-mediated immune responses (IBMIR) in NK cells**. Dimensional reduction analysis depicting cell subpopulation clusters for (**A**) control blood samples, (**B**) fresh and vitrified/rewarmed (VR) islets, and (**C**) fresh and VR SC-islets. (**D**) Transformed relative expression of key cell surface markers identified in A-C. Population sizes for control blood samples, fresh islets/ SC-islets, and VR islets/SC-islets are shown for each cluster in (**E**) islets and (**F**) SC-islets. Data are mean ± 95% confidence interval. Bubble plots illustrate the relative population size compared to control for (**G**) fresh and VR islets and (**H**) fresh and VR SC-islets.

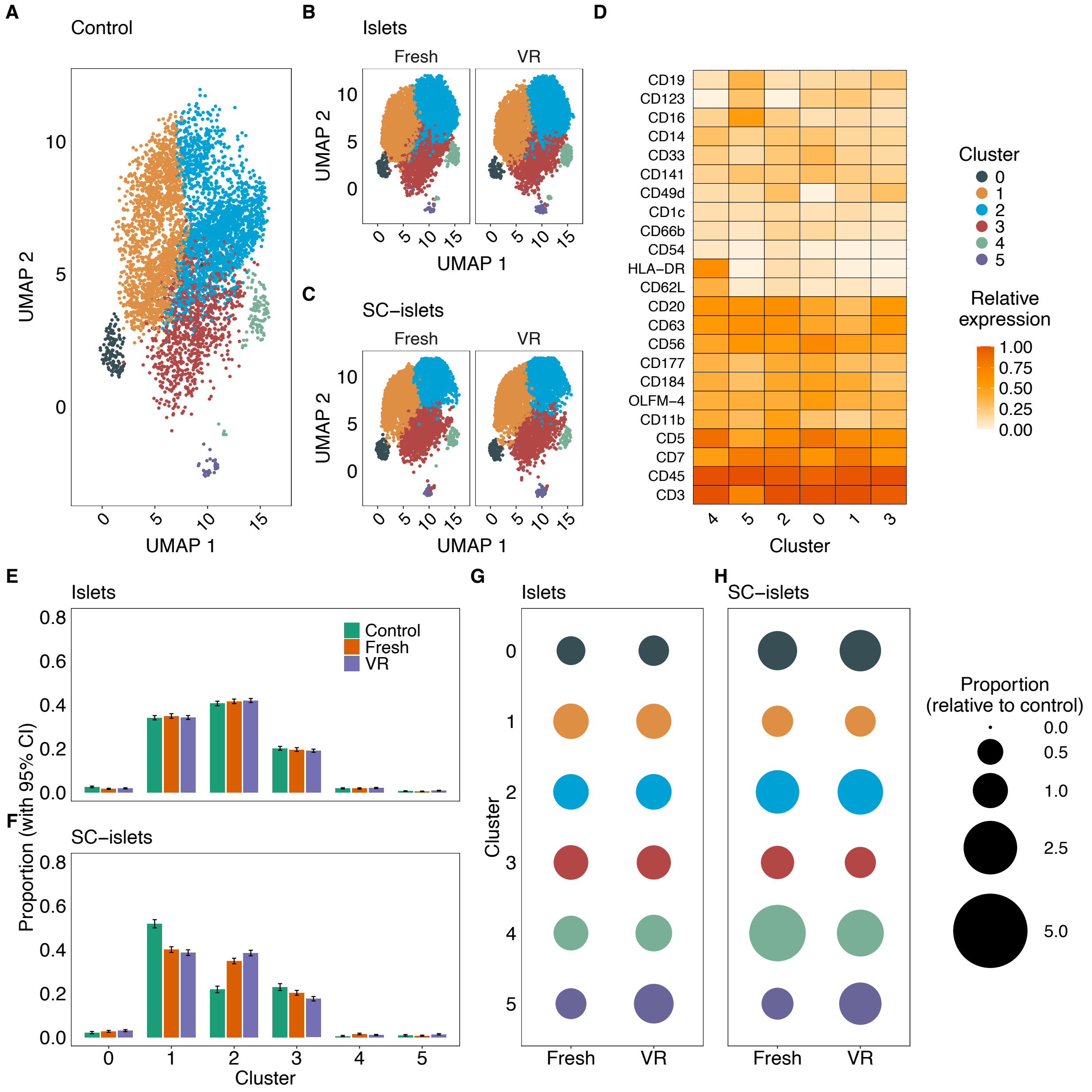


**Fig. S18. Instant blood-mediated immune responses (IBMIR) in NKT cells**. Dimensional reduction analysis depicting cell subpopulation clusters for (**A**) control blood samples, (**B**) fresh and vitrified/rewarmed (VR) islets, and (**C**) fresh and VR SC-islets. (**D**) Transformed relative expression of key cell surface markers identified in A-C. Population sizes for control blood samples, fresh islets/ SC-islets, and VR islets/SC-islets are shown for each cluster in (**E**) islets and (**F**) SC-islets. Data are mean ± 95% confidence interval. Bubble plots illustrate the relative population size compared to control for (**G**) fresh and VR islets and (**H**) fresh and VR SC-islets.

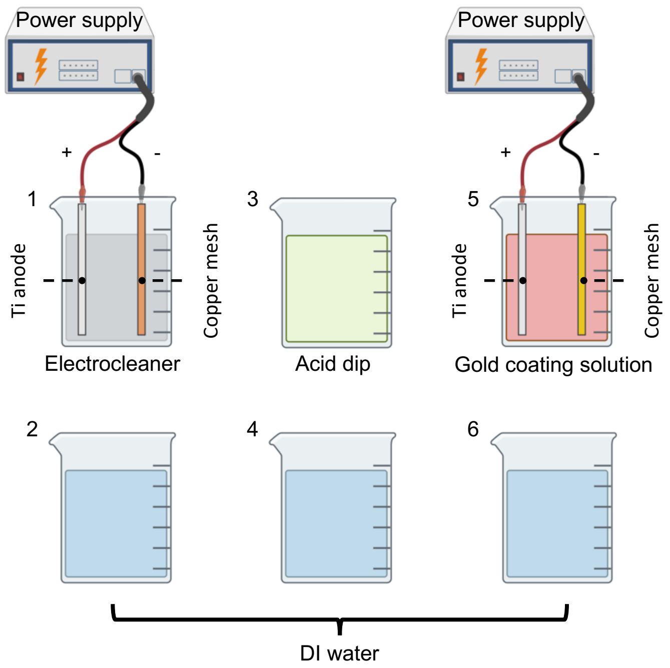


**Fig. S18. Schematic of the gold electroplating process used to fabricate copper–gold CryoMesh.** The process involves six steps: (1) electrocleaning; (2) rinse with deionized (DI) water; (3) acid dip; (4) rinse with DI water; (5) gold electroplating; and (6) rinse with DI water. A final acid dip and DI water rinse are performed after step 6. DI, deionized water.

### Supplementary Tables

#### Table S1. Characteristics of human pancreas donors and islet isolations.

| | Donor 1 | Donor 2 | Donor 3 | Donor 4 |
|---|---|---|---|---|
| **Age (y)** | 31 | 52 | 46 | 35 |
| **Sex** | F | F | F | F |
| **Ethnicity** | Hispanic | Caucasian | Caucasian | Caucasian |
| **BMI** | 52.35 | 43.32 | 33.95 | 45.96 |
| **Cause of Death** | Anoxia | CVS/ Stroke | CVS/ Stroke | Anoxia |
| **Serum Amylase (U/l)** | 50 | 21 | 94 | 39 |
| **Serum Lipase (U/l)** | 52 | 23 | 16 | 6 |
| **Serum Creatinine (mg/dl)** | 0.35 | 0.82 | 0.7 | 1.09 |
| **Cold Ischemia Time** | 5h 0min | 5h 3min | 6h 10min | 5h 52min |
| **Pancreas Weight (g)** | 118.9 | 69.4 | 93 | 91 |
| **Digestion Time** | 37 min | 42 min | 46 min | 39 min |
| **Culture Time** | 42 hours | 46 hours | 24 hours | 48 hours |
| **Purity** | 90% | 80% | >95% | 85% |
| **Embedded** | 40% | 30% | <5% | 20% |
| **Fragmented** | 1% | 1% | 5% | 1% |
| **IEQ (purified)** | 675,254 | 635,280 | 554,489 | 737,322 |
| **IEQ (VR)** | 462,655 | 351,033 | 412,696 | 585,361 |

**Table S2. Cryopreservation efficiency of pancreatic islet clusters.**

| Method | Islet cluster per loading | Cooling processing time (mins) | Rewarming processing time (mins) | Viability (%) | Target number† | Total time (h) | Sophisticated equipment and training |
|---|---|---|---|---|---|---|---|
| Manual laser rewarming process (*23, 79*) | 15* | > 3 | > 3 | 62% | 1,000,000 | > 6,666 | Yes |
| Idealized automated laser rewarming process | 15* | > 0.5 | > 0.5 | 62% | 1,000,000 | > 1,111 | Yes |
| Nylon mesh (*23*) | 4,000** | < 2 | < 2 | 92% | 1,000,000 | < 16.6 | No |
| CryoMesh (This work) | 300,000-600,000*** | < 2 | < 2 | 92% | 1,000,000 | < 0.22 | No |

† Target number refers to the number of pancreatic islet clusters after cryopreservation.
* The number to achieve the highest direct post-rewarming viability is around 62%, which might not be able to be used for further transplant.
** The number is based on the theoretical density of a monolayer islet cluster on the nylon mesh (Fig. 2B).
*** The number is based on a 7x4 cm copper-gold mesh (larger mesh sizes are possible).

**Table S3. Top 20 differentially expressed genes between fresh and vitrified/rewarmed (VR) native pancreatic islets identified by RNA-seq.**

| Gene Name | $log_2FC$ | $log_2CPM$ | F | P Value | FDR | Direction |
|---|---|---|---|---|---|---|
| GMPR2 | -1.847 | 1.239 | 40.46 | $2.11 \times 10^{-4}$ | 0.4372 | DOWN |
| GSDMD | 2.030 | 0.238 | 31.13 | $5.16 \times 10^{-4}$ | 0.4372 | UP |
| INTS3 | 1.569 | 1.125 | 30.08 | $5.70 \times 10^{-4}$ | 0.4372 | UP |
| DCAF11 | 1.522 | 1.166 | 29.38 | $6.14 \times 10^{-4}$ | 0.4372 | UP |
| FOSB | -2.822 | 5.443 | 26.02 | $9.06 \times 10^{-4}$ | 0.4372 | DOWN |
| MIR221 | -1.787 | -1.776 | 24.97 | $1.04 \times 10^{-3}$ | 0.4372 | DOWN |
| IFI27 | -1.794 | -0.967 | 23.49 | $1.26 \times 10^{-3}$ | 0.4372 | DOWN |
| LINC00520 | 2.103 | -1.007 | 22.53 | $1.29 \times 10^{-3}$ | 0.4372 | UP |
| ENSG00000287689 | -2.967 | -0.518 | 20.72 | $1.41 \times 10^{-3}$ | 0.4372 | DOWN |
| YTHDC1 | -1.030 | 1.431 | 22.25 | $1.47 \times 10^{-3}$ | 0.4372 | DOWN |
| RPL7AP43 | -1.875 | -1.135 | 22.32 | $1.49 \times 10^{-3}$ | 0.4372 | DOWN |
| VNN3P | 1.208 | 0.402 | 21.62 | $1.61 \times 10^{-3}$ | 0.4372 | UP |
| LINC02635 | -2.032 | -0.781 | 20.92 | $1.65 \times 10^{-3}$ | 0.4372 | DOWN |
| FGF2 | 0.696 | 6.756 | 21.16 | $1.72 \times 10^{-3}$ | 0.4372 | NO |
| ZNF536 | -1.604 | -1.562 | 20.74 | $1.86 \times 10^{-3}$ | 0.4372 | DOWN |
| ENSG00000228719 | -1.570 | -1.486 | 20.43 | $1.91 \times 10^{-3}$ | 0.4372 | DOWN |
| MYADM | -0.731 | 6.025 | 20.16 | $1.99 \times 10^{-3}$ | 0.4372 | NO |
| LINC02029 | -1.264 | -0.093 | 19.97 | $2.05 \times 10^{-3}$ | 0.4372 | DOWN |
| SLC22A18 | 1.307 | -0.505 | 19.88 | $2.10 \times 10^{-3}$ | 0.4372 | UP |
| ENSG00000255434 | 1.242 | -0.661 | 19.73 | $2.15 \times 10^{-3}$ | 0.4372 | UP |

CPM, counts per million reads; F, F statistic; FC, fold change; FDR, false discovery rate

**Table S4. Top 20 differentially expressed genes between fresh and vitrified/rewarmed (VR) SC-islets identified by RNA-seq.**

| Gene Name | $log_2FC$ | $log_2CPM$ | F | P Value | FDR | Direction |
|---|---|---|---|---|---|---|
| PPP1R15A | 2.175 | 7.056 | 385.49 | $1.82 \times 10^{-6}$ | 0.0221 | UP |
| KLKP1 | 5.343 | -1.453 | 101.00 | $2.07 \times 10^{-6}$ | 0.0221 | UP |
| GADD45A | 1.679 | 5.154 | 183.87 | $1.45 \times 10^{-5}$ | 0.0699 | UP |
| LINC00511 | 1.763 | 4.534 | 181.65 | $1.50 \times 10^{-5}$ | 0.0699 | UP |
| HSPA7 | 3.580 | -0.449 | 119.47 | $2.03 \times 10^{-5}$ | 0.0699 | UP |
| RFPL3S | 2.337 | 2.335 | 147.80 | $2.67 \times 10^{-5}$ | 0.0699 | UP |
| ADM | 2.372 | 3.421 | 144.43 | $2.84 \times 10^{-5}$ | 0.0699 | UP |
| ENSG00000261889 | 3.085 | -1.417 | 111.45 | $3.05 \times 10^{-5}$ | 0.0699 | UP |
| INHBA | 1.969 | 5.099 | 139.48 | $3.13 \times 10^{-5}$ | 0.0699 | UP |
| TNFRSF10D | 1.478 | 4.848 | 136.50 | $3.32 \times 10^{-5}$ | 0.0699 | UP |
| KLF6 | 1.588 | 7.543 | 131.86 | $3.65 \times 10^{-5}$ | 0.0699 | UP |
| SQSTM1 | 2.483 | 3.834 | 126.91 | $4.06 \times 10^{-5}$ | 0.0699 | UP |
| NFKBIZ | 1.618 | 6.433 | 122.56 | $4.47 \times 10^{-5}$ | 0.0699 | UP |
| ATF3 | 2.733 | 6.522 | 121.39 | $4.59 \times 10^{-5}$ | 0.0699 | UP |
| HSPD1P10 | 3.102 | -1.111 | 104.07 | $5.30 \times 10^{-5}$ | 0.0701 | UP |
| RND1 | 1.757 | 4.769 | 113.78 | $5.48 \times 10^{-5}$ | 0.0701 | UP |
| VMP1 | 1.169 | 7.393 | 108.82 | $6.19 \times 10^{-5}$ | 0.0719 | UP |
| TNFAIP3 | 1.628 | 5.724 | 107.51 | $6.40 \times 10^{-5}$ | 0.0719 | UP |
| ADGRF2 | 2.660 | -1.334 | 84.15 | $7.11 \times 10^{-5}$ | 0.0724 | UP |
| RASGRP3 | 2.683 | 2.700 | 103.42 | $7.12 \times 10^{-5}$ | 0.0724 | UP |

CPM, counts per million reads; F, F statistic; FC, fold change; FDR, false discovery rate

**Table S5. Function of top 20 differentially expressed genes between fresh and vitrified/rewarmed (VR) SC-islets identified by RNA-seq.**

| Gene Name | Function / Role Summary | Notes |
|---|---|---|
| PPP1R15A | Regulates stress responses by directing dephosphorylation of eIF2α; plays a role in the unfolded protein response and apoptosis (*70*). | Also known as GADD34. |
| KLKP1 | Limited characterization; potentially a pseudogene or uncharacterized locus (*80*). | Functional details are sparse in the literature. |
| GADD45A | Involved in DNA damage response, cell cycle arrest, and apoptosis; mediates growth arrest after stress or genotoxic damage (*81*). | Part of the GADD45 family responding to various stress signals. |
| LINC00511 | A long intergenic non-coding RNA implicated in the regulation of gene expression; associated with cancer progression and cellular proliferation (*82*). | Non-coding RNA – function is regulatory rather than enzymatic. |
| HSPA7 | Member of the Hsp70 family; likely functions as a molecular chaperone assisting in protein folding during cellular stress responses (*72*). | Although similar to other heat shock proteins, its precise role is less well characterized. |
| RFPL3S | Poorly characterized; may be related to the RING finger protein family suggesting a potential role in ubiquitination or proteasomal degradation pathways (*80*). | Detailed functional information is limited. |
| ADM | Encodes adrenomedullin, a peptide involved in vasodilation, angiogenesis, and regulation of hormone secretion; plays roles in cardiovascular and metabolic homeostasis (*83*). | Acts as both a hormone and a paracrine factor. |
| ENSG00000261889 | An uncharacterized transcript (often a long non-coding RNA) with unknown or predicted regulatory functions. | |
| INHBA | Encodes the inhibin beta A subunit; part of the transforming growth factor-beta (TGF-β) superfamily; involved in regulating cell proliferation, differentiation, and reproductive biology (*84*). | Forms activin and inhibin complexes depending on the context. |
| TNFRSF10D | Functions as a decoy receptor for TRAIL (TNF-related apoptosis-inducing ligand); modulates apoptosis by preventing the activation of death signaling pathways (*73*). | Its decoy activity helps protect cells from TRAIL-induced apoptosis. |
| KLF6 | A transcription factor involved in regulating cell differentiation, proliferation, and tumor suppression; influences various gene expression programs (*85*). | Often studied in the context of cancer as a potential tumor suppressor. |
| SQSTM1 | Acts as an adaptor protein in autophagy and mediates signaling pathways including NF-κB; involved in protein turnover and cellular stress responses (*74*). | Also known as p62; mutations are linked to neurodegeneration and other diseases. |
| NFKBIZ | Regulates inflammatory responses by modulating NF-κB activity; functions as a nuclear inhibitor that can fine-tune immune and stress response gene expression (*75*). | Also referred to as IκBζ. |
| ATF3 | A stress-induced transcription factor that modulates gene expression in response to various cellular stresses, including inflammation and DNA damage (*76*). | Acts as both a transcriptional activator and repressor. |
| HSPD1P10 | Likely a pseudogene related to the mitochondrial chaperonin HSPD1; its transcriptional activity and functional impact, if any, are not well established (*80*). | Pseudogene |
| RND1 | A member of the Rho GTPase family that regulates cytoskeletal organization and cell migration; involved in modulating cell shape and motility (*86*). | Unlike typical Rho GTPases, RND1 is constitutively active due to its low GTPase activity. |

| | | |
|---|---|---|
| VMP1 | Plays a key role in autophagy and the formation of autophagosomes; also implicated in processes like cell adhesion and membrane trafficking (*87*). | Involved in cellular homeostasis through its role in autophagic pathways. |
| TNFAIP3 | Encodes a ubiquitin-editing enzyme that serves as a negative regulator of NF-κB signaling; helps limit inflammatory responses and cell survival signals (*77*). | Also known as A20; mutations are linked to autoimmunity and lymphomas. |
| ADGRF2 | Possible pseudogene. Predicted to be involved in adenylate cyclase-activating G protein-coupled receptor signaling pathway (*80*). | Also known as GPR111. Likely pseudogene |
| RASGRP3 | Functions as a guanine nucleotide exchange factor (GEF) for Ras family GTPases; plays a role in activating Ras signaling pathways which regulate cell proliferation and differentiation (*88*). | Implicated in immune cell signaling and vascular development. |

**Table S6. Cluster phenotypes and activation status for neutrophils.**

| Cluster | Phenotype | Key Markers (Relative) | Interpretation |
|---|---|---|---|
| 0 | Resting / Quiescent | $CD11b^{neg}$, $CD62L^{low/mid}$ | Baseline: quiescent. |
| 1 | Resting | $CD11b^{low}$, $CD62L^{mid}$ | Baseline: Similar to 0. |
| 2 | Resting | $CD11b^{low}$, $CD62L^{high}$ | Baseline: High L-selectin indicates a classical resting state. |
| 3 | Transitioning | $CD11b^{mid}$, $CD62L^{high}$ | Slight upregulation of CD11b compared to 0-2. |
| 4 | CD177+ Subset | $CD177^{high}$ | Distinct Subset: Represents the CD177+ population (approx. 50% of donor cells). |
| 5 | Aged / Senescent | $CD184^{high}$, $CD62L^{neg}$ | Distinct Fate: High CXCR4 indicates aging/homing to bone marrow. A divergent path from acute activation. |
| 6 | Primed | $CD11b^{high}$, $CD62L^{high}$ | Mobilization: Integrins are upregulated, but L-selectin is retained. |
| 7 | Strongly Primed | $CD62L^{max}$, $CD11b^{high}$ | Peak Priming: Maximal retention of L-selectin despite high integrin expression. |
| 8 | Activated (Early) | $CD11b^{high}$, $CD62L^{low}$ | Loss of L-selectin (CD62L) begins, marking the transition to full activation. |
| 9 | Activated | $CD11b^{high}$, $CD66b^{high}$ | Consistent activation profile. |
| 10 | Hyper-Adhesive | $CD11b^{max}$, $CD54^{max}$ | Peak Adhesion: Highest expression of adhesion molecules (Mac-1, ICAM-1). |
| 11 | Degranulating | $CD66b^{max}$, $CD62L^{low}$ | Peak Effector: Maximal degranulation marker expression. Represents the endpoint of the activation cascade. |

**Table S7. Cluster phenotypes for monocytes, dendritic cells, and myeloid-derived suppressor cells.**

| Cluster | Phenotype | Informative Markers | Activation Markers | Activation Status |
|---|---|---|---|---|
| 0 | Non-Classical Monocyte | $CD16^{high}$, $CD14^{low}$, $CD62L^{neg}$ | $CD16^{pos}$, $CD62L^{neg}$ | Patrolling |
| 1 | Intermediate Monocyte | $CD177^{high}$, $CD56^{high}$, $CD16^{mid}$ | $CD177^{high}$, $CD56^{high}$ | Transitioning |
| 2 | Classical Monocyte | $CD14^{high}$, $CD33^{high}$, $CD16^{low}$ | $CD11b^{high}$, HLA-DR+ | Inflammatory |
| 3 | Classical Monocyte (Activated) | $CD14^{high}$, $CD63^{high}$, $CD11b^{high}$ | $CD63^{high}$ (Degranulation), $CD11b^{high}$ | Highly Activated |
| 4 | pDC (Quiescent) | $CD123^{high}$, HLA-$DR^{mid}$ | HLA-$DR^{low}$, No Degranulation | Quiescent |
| 5 | pDC (Mature) | HLA-$DR^{high}$, $CD1c^{high}$, $CD33^{high}$ | HLA-$DR^{high}$, $CD1c^{high}$ | Mature / Antigen Presenting |
| 6 | G-MDSC | $CD66b^{high}$, $CD54^{high}$, HLA-$DR^{neg}$ | $CD54^{high}$ (ICAM-1), $CD66b^{high}$ | Suppressive |

**Table S8. Cluster phenotypes for NK cell gate.**

| Cluster | High Markers | Low Markers | Activation Markers | Activation Status |
|---|---|---|---|---|
| 0 | CD177, CD7 | CD123, CD14, CD16, CD19, CD1c, CD20, CD5, CD54, CD56, OLFM-4 | CD66b↑, CD11b↑, CD62L↓ | Moderate |
| 1 | CD123, CD14, CD141, CD184, CD3, CD5 | CD177, CD19, CD20, CD45, CD63, CD7 | CD11b↑, OLFM-4↑ | Moderate |
| 2 | CD56 | CD141, CD184, CD3, CD66b, HLA-DR | CD63↑ | Moderate |
| 3 | CD184, CD19, CD20, CD3, CD66b, HLA-DR | CD11b, CD123, CD14, CD141, CD16, CD49d, CD5, CD54, CD63, OLFM-4 | CD66b↑, CD62L↓ | Moderate |
| 4 | CD16, CD3, CD33, CD45, CD66b | CD11b, CD1c, CD56, CD62L | CD66b↑, OLFM-4↑, CD62L↓ | High |
| 5 | CD16, CD177, CD45, OLFM-4 | CD33, CD49d, CD62L | CD63↑, CD66b↑, OLFM-4↑, CD62L↓ | High |
| 6 | CD11b, CD19, CD1c, CD20, CD49d, CD54, CD56, CD62L, CD63, CD7 | CD184, CD3, CD33, HLA-DR | CD63↑, CD11b↑ | High |
| 7 | CD11b, CD123, CD14, CD141, CD1c, CD33, CD49d, CD5, CD54, CD62L, CD63, HLA-DR, OLFM-4 | CD177, CD45, CD66b, CD7 | CD63↑, CD11b↑, CD54↑, OLFM-4↑ | High |

### Table S9. Cluster phenotypes for NK-T cell gate.

| Cluster | High Markers | Low Markers | Activation Markers | Activation Status |
|---|---|---|---|---|
| 0 | CD33, CD66b, HLA-DR | CD11b, CD14, CD16, CD177, CD1c, CD20, CD3, CD45, CD49d, CD5, CD54, CD62L, CD63, CD7 | CD66b↑, CD62L↓ | Moderate |
| 1 | CD123, CD141, CD184, CD1c, CD3, CD5, CD56, CD62L, CD7, OLFM-4 | CD11b, CD14, CD20, CD54, CD63, CD66b, HLA-DR | OLFM-4↑, CD62L↓ | Moderate |
| 2 | CD11b, CD177, CD184, CD20, CD3, CD45, CD49d, CD63, CD7 | CD123, CD141, CD19, CD1c, CD33, CD62L, OLFM-4 | CD63↑, CD11b↑, CD62L↓ | High |
| 3 | CD19, CD45, CD49d | CD16, CD184, CD33, CD56, HLA-DR | CD63↑, OLFM-4↑, CD62L↓ | High |
| 4 | CD11b, CD14, CD16, CD177, CD33, CD45, CD5, CD54, CD62L, CD66b, HLA-DR | CD123, CD141, CD19, CD49d, CD56, CD7, OLFM-4 | CD63↑, CD66b↑, CD11b↑, CD54↑ | High |
| 5 | CD123, CD14, CD141, CD16, CD19, CD1c, CD20, CD54, CD56, CD63, OLFM-4 | CD177, CD184, CD3, CD33, CD45, CD5, CD66b, HLA-DR | CD63↑, CD54↑, OLFM-4↑, CD62L↓ | High |
